# Multiplexed *in vivo* base editing identifies functional gene-variant-context interactions

**DOI:** 10.1101/2025.02.23.639770

**Authors:** Jonuelle Acosta, Grace A. Johnson, Samuel I. Gould, Kexin Dong, Yovel Lendner, Diego Detrés, Ondine Atwa, Jari Bulkens, Samuel Gruber, Manuel E. Contreras, Alexandra N. Wuest, Varun K. Narendra, Michael T. Hemann, Francisco J. Sánchez-Rivera

## Abstract

Human genome sequencing efforts in healthy and diseased individuals continue to identify a broad spectrum of genetic variants associated with predisposition, progression, and therapeutic outcomes for diseases like cancer^1–6^. Insights derived from these studies have significant potential to guide clinical diagnoses and treatment decisions; however, the relative importance and functional impact of most genetic variants remain poorly understood. Precision genome editing technologies like base and prime editing can be used to systematically engineer and interrogate diverse types of endogenous genetic variants in their native context^7–9^. We and others have recently developed and applied scalable sensor-based screening approaches to engineer and measure the phenotypes produced by thousands of endogenous mutations *in vitro*^10–12^. However, the impact of most genetic variants in the physiological *in vivo* setting, including contextual differences depending on the tissue or microenvironment, remains unexplored. Here, we integrate new cross-species base editing sensor libraries with syngeneic cancer mouse models to develop a multiplexed *in vivo* platform for systematic functional analysis of endogenous genetic variants in primary and disseminated malignancies. We used this platform to screen 13,840 guide RNAs designed to engineer 7,783 human cancer-associated mutations mapping to 489 endogenous protein-coding genes, allowing us to construct a rich compendium of putative functional interactions between genes, mutations, and physiological contexts. Our findings suggest that the physiological *in vivo* environment and cellular organotropism are important contextual determinants of specific gene-variant phenotypes. We also show that many mutations and their *in vivo* effects fail to be detected with standard CRISPR-Cas9 nuclease approaches and often produce discordant phenotypes, potentially due to site-specific amino acid selection- or separation-of-function mechanisms. This versatile platform could be deployed to investigate how genetic variation impacts diverse *in vivo* phenotypes associated with cancer and other genetic diseases, as well as identify new potential therapeutic avenues to treat human disease.

## Main

Diverse types of germline and somatic genetic alterations have been linked to the development, progression, and therapeutic outcomes of cancer and other human diseases^13,14^. In some cases, specific mutant genes or genetic variants have been shown to drive or influence the course of a given disease^15–17^, serving as potential diagnostic or therapeutic targets. These include single nucleotide variants (SNVs) in oncogenes like *KRAS*, *BRAF*, and *EGFR*, among others, which have been extensively profiled and known to be associated with distinct oncogenic potential and therapeutic sensitivity^18–21^. More broadly, molecular profiling of human cancer cell lines and tumor biopsies using next-generation DNA sequencing technologies has led to the identification of a broad spectrum of complex genetic alterations that may promote or otherwise influence cancer development, evolution, and therapy responses^1–6^. These technologies are quickly becoming a mainstay in the clinic and are already influencing disease diagnosis and clinical decision-making in real time^22^, highlighting the urgent need to assess the functional impact of each of these variants in disease pathogenesis.

Most human disease-associated mutations remain classified as variants of unknown significance (VUS)^23^. Efforts to functionally examine disease-associated variants at scale have been challenging in large part due to technical limitations. Recent developments in precision genome editing methods, such as base editing^7,8^ and prime editing^9^, overcome many of the limitations of previous technologies by allowing the systematic engineering and interrogation of diverse types of genetic variants in their native endogenous contexts^24^. Base editors are comprised of programmable Cas9 nickases fused to cytosine or adenosine deaminase domains^25^. These editors can be used to engineer SNVs, including transitions (e.g. C•G to T•A or A•T to G•C) and, more recently, transversions (e.g. A•T to C•G) upon being targeted to endogenous loci by single guide RNAs (sgRNAs)^7,8,26–28^. Base editing technology enables the modeling of clinically relevant genetic alterations that were previously difficult to examine with standard Cas9 nuclease technology^29–31^. Base and prime editing are also scalable technologies; indeed, recent studies have shown that high-throughput *in vitro* base and prime editing screens can be used for comprehensive functional investigation of variants within a single gene, or across a broad range of genetic loci^10–12,32–45^.

*In vitro* tissue culture studies have been instrumental in elucidating many fundamental principles of biology and disease. However, the physiological effects of diverse types of genetic and pharmacological perturbations are not always recapitulated in standard *in vitro* culture conditions and, in some cases, can be the exact opposite of autochthonous *in vivo* phenotypes^46–63^. Physiological variables like nutrient availability^64,65^, oxygen concentration^66,67^, heterotypic cell-cell interactions in native tissue niches^68–70^, proximity to the circulation^71–73^, drug penetration/availability^74,75^, and the presence or absence of an immune system^76^, among others, have been difficult to model in cell culture settings, highlighting the need to accurately recapitulate autochthonous physiological conditions when assessing the effects of genetic perturbations.

A number of obstacles have prevented the deployment of high-throughput base editing technologies *in vivo* to understand how endogenous genes and mutations differentially impact the biology of cells within distinct tissue microenvironments or promote systemic phenotypes like metastatic dissemination and organotropism. These include (1) computational challenges related to the design of cross-species sgRNA libraries to accurately engineer evolutionarily conserved mutations, (2) the paucity of robust immunocompetent models capable of accurately representing ultracomplex sgRNA libraries designed to model diverse types of mutations, (3) fitness and immunogenicity issues related to stable expression of Cas9-based cytosine (CBE) and adenosine (ABE) base editors^77,78^, and (4) the lack of integrative methods that allow for simultaneous empirical measurement of dynamic sgRNA editing activities in cells or tissues of interest coupled to functional purification of cells that exhibit the desired editing properties, among others. To address the first of these challenges, we recently developed a versatile computational method called H2M (https://human2mouse.com/) that can be used for high-throughput *in silico* modeling of most types of human mutations, as well as for designing variant-specific cross-species reagents and libraries that can be used to systematically model and interrogate equivalent mutations across species (Dong et al., *in revision*).

In this study, we attempted to overcome the remaining three challenges by integrating ‘hit-and-run’ base editing with new cross-species base editing sensor libraries, editing activity fluorescent reporters, and syngeneic mouse models to develop a scalable *in vivo* platform for high-throughput functional analysis of genetic variants in animals harboring an intact immune system. We used this platform to screen 13,840 H2M-designed sgRNAs *in vivo* to investigate the functional impact of 7,783 functionally diverse human cancer-associated mutations mapping to 489 endogenous protein-coding mouse genes. We show that this platform can be used to identify *bona fide* oncogenic and tumor suppressor gene mutations associated with diverse types of human cancers, as well as functionally-distinct mutations mapping to the same gene/protein that are predicted to cause different biochemical and biophysical activities. Moreover, parallel *in vitro* and *in vivo* screens suggest that the physiological *in vivo* environment is an important contextual determinant of variant-specific phenotypes, including cellular organotropism and meningeal infiltration. Importantly, we also show that many mutations and their related phenotypes fail to be detected with traditional CRISPR-Cas9 nuclease approaches and, in some cases, produce discordant functional phenotypes potentially due to site-specific amino acid selection- or separation-of-function molecular mechanisms. This versatile platform could be readily deployed to systematically investigate how genetic variation impacts *in vivo* phenotypes associated with cancer and other genetic diseases, as well as identify new genotype-specific therapeutic strategies to improve patient outcomes.

## Results

### Next-generation base editing libraries to model human genetic variation

We recently developed H2M (human-to-mouse) (https://human2mouse.com/), a new computational pipeline to analyze human genetic variation data and systematically model and predict the functional consequences of equivalent mouse variants (Dong et al., *in revision*). We implemented H2M and PEGG (pegg.base module)^11,79^ to design a new cross-species base editing sensor library to comprehensively model human cancer-associated somatic variants identified in MSK-IMPACT patient cohorts^1^ (**Fig. 1a**). Clinical genomic data collected from >61,000 tumor samples was filtered for genetic variants that were present in ≥ 3 patients and amenable to adenine (ABE) and cytosine (CBE) base editing (**Fig. 1b**). Out of 19,020 amenable variants, H2M mapped 10,948 variants to the mouse genome that contained ≥ 2 homologous amino acids proximal to the targeted amino acid (flank size ≥ 2) **(Extended Data Fig. 1a)**. In total, we identified 7,783 somatic variants that could be modeled in mouse cells using flexible Cas9 nickases that recognize NG protospacer adjacent motifs (PAMs)^80–82^. In addition, we only included sgRNAs where the target base fell within the +4 to +8 region of the protospacer, which was previously shown to be the region that exhibits maximal base editing (optimal editing window) (**Extended Data Fig. 1b)**^81,82^. Our computational pipeline identified an average of 1.65 sgRNAs per variant across a total of 489 genes **(Fig. 1c, Extended Data Fig. 1c-d).** The final library (MBESv2) contained 13,840 sgRNAs targeting 7,783 unique mutations, including multiple highly represented genes targeted by >100 independent sgRNAs designed to engineer cancer-associated mutations **(Fig. 1c).** This library also included 581 ‘legacy’ sgRNAs from MBESv1^7,8,10^ that did not pass stringent H2M mapping and/or homology thresholds, but were included to enable comparison between MBESv1 and v2 libraries (**Fig. 1d**). Consistent with the distribution of SNVs from the clinical genomic data, the MBESv2 library is largely composed of CBE sgRNAs (n=12,806); thus, we sub-pooled the library by base editing type to screen A•T to G•C (ABE) and C•G to T•A (CBE) mutations separately to minimize experimental noise (**Fig. 1d**, **Extended Data Fig. 1e-f)**. Importantly, all sgRNAs were paired with variant-specific synthetic ‘sensor’ sites that have been shown to accurately recapitulate editing patterns observed at their cognate endogenous target locus^10,11^ (**Fig. 1e, Extended Data Fig. 1g**). We and others have recently shown that sensor-based readouts can be coupled with high-throughput multiplexed genetic screens to quantitatively and dynamically assess sgRNA editing patterns at cognate endogenous loci, enhancing experimental robustness and streamlining downstream analyses^10,11^.

**Figure 1.**
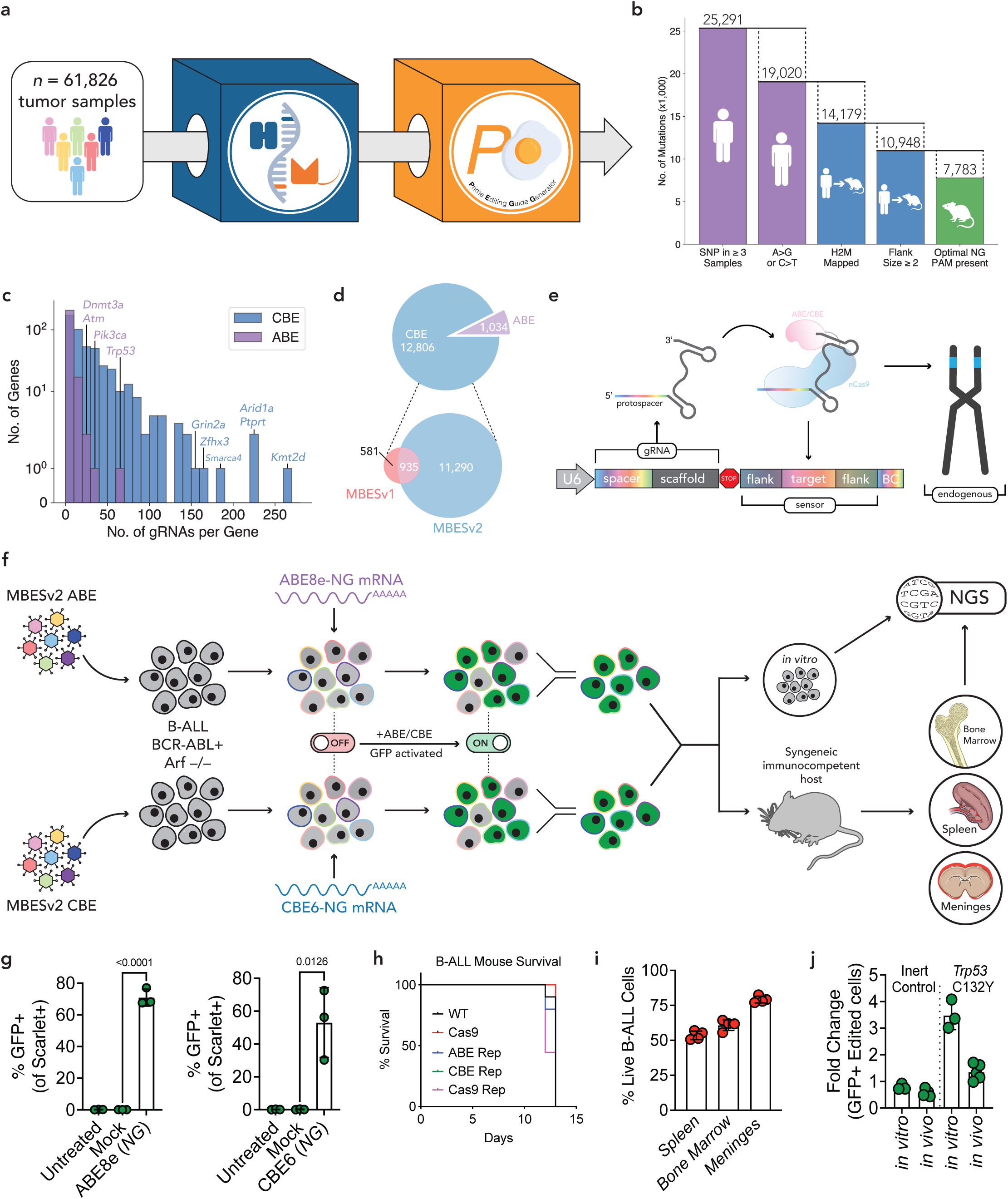
Development of next-generation base editing libraries to accurately model human genetic variation in immunocompetent mice. **a,** Schematic of library generation process. Base editing amenable SNPs from 61,826 tumor samples were filtered for appearance in ≥3 samples. This mutation set was mapped to the mouse genome with H2M (Dong et al., *in revision*), filtered to have a flank size ≥2, and then fed into PEGG^11^ to generate an NG PAM base editing gRNA library. **b,** Barplot showing the number of mutations remaining after each filtration step. **c,** Histogram of the number of gRNAs per gene for both the CBE and ABE libraries. Selected highly represented genes labeled. **d,** Pie chart of ABE and CBE targeting guides (top) and venn diagram showing the overlap between MBESv1 (refs. 7,8,10) and MBESv2 included in the CBE subpool. **e,** Schematic of the base editing sensor construct. BC, barcode. **f,** Overall schematic of screening approach. B-ALL cells are transduced with MBES libraries (ABE or CBE) at low MOI, before electroporation with base editor mRNA. Cells are sorted for base editor reporter GFP activation, and then delivered to mice or kept *in vitro*. Tissue samples are then harvested and prepared for NGS. **g,** Flow-cytometry assisted quantification of % GFP positive ABE or CBE reporter cells after 48hrs of electroporation with *in vitro* transcribed mRNAs encoding ABE8e-NG or CBE6-NG. Each symbol represents a biological replicate from n=3 independent experiments. Error bars represent mean ± s.d. Statistical significance determined by two-tailed Student’s *t*-test **h,** Kaplan-Meir survival analysis of C57BL/6J mice transplanted with WT, ABE, and CBE reporter B-ALL cells via tail vein injection. **i,** Flow-cytometry assisted quantification of mScarlet+ CBE reporter cells in the spleen, bone marrow, and meninges of leukemic mice t=10 days post-transplantation. **j,** Side-side comparison of tandem *in vitro* and *in vivo* competition assays of B-ALL Trp53 WT (inert control) and *Trp53* C135Y mutant cells. 1% of GFP+ *Trp53* WT *or* C135Y mutant cells were mixed with unlabeled controls at t=0 and differences in growth were measured after 10 days by quantifying the fold-change of GFP+ cells using flow cytometry. See also Extended Figures 1-3.

### Hit-and-run base editing allows multiplexed *in vivo* genetic analysis in immunocompetent mice

Our groups have previously shown that syngeneic mouse models of B cell lymphoblastic acute leukemia (B-ALL) driven by the oncogenic *BCR-ABL* fusion (hereafter *BCR-ABL+* B-ALL)^83^ can be used for high content *in vivo* genetic screens^50,56,57^. The advantages of this model are three-fold. First, cells derived from this model have a very high leukemia-initiating potential, allowing us to introduce and robustly examine thousands of unique genetic perturbations even within single animals. Second, these cells can be orthotopically transplanted into syngeneic, immunocompetent C57BL/6J hosts to closely recapitulate *in vivo* physiological conditions. Lastly, transplanted B-ALL cells faithfully recapitulate the clinical progression and dissemination of human leukemia into the spleen, bone marrow, and meninges, allowing us to examine if and how genes or specific mutations influence leukemia organ-specific growth and organotropism^83,84^.

To develop a multiplexed *BCR-ABL+* B-ALL screening platform to investigate the impact of cancer-associated mutations *in vivo*, we incorporated quantitative base editing activity reporters into B-ALL cells^85^ (**Figure 1f**). These reporters toggle GFP expression ON only after a productive base editing event enables the transcription and translation of a GFP mRNA transcript^85^ (**Extended Data Fig. 2a**). In the course of these experiments, we noted that B-ALL cells were highly sensitive to stable expression of ABEs and CBEs (**Extended Fig. 2b-d**), consistent with recent work showing that these enzymes can sometimes induce genotoxic effects in cells from the hematopoietic system^86^. Combined with the well-established observation that stable expression of Cas9 proteins can be strongly immunogenic *in vivo*, we tested whether transient ‘hit-and-run’ base editing using mRNA could avoid the potential cellular toxicity and immunogenicity associated with base editors. To do so, we electroporated base editing reporter-expressing cells with optimized *in vitro* transcribed mRNAs encoding ABE8e or CBE6^81,82^. This ‘hit and run’ approach resulted in high base editing activity at both ABE and CBE reporters, causing cells to become GFP+ and remain viable throughout the course of the experiment (**Fig. 1g**). We also confirmed that reporter expression did not influence disease latency, penetrance, or leukemia cell occupancy across any of the tissues and organs we examined in leukemia-bearing mice (**Fig. 1h**).

Next, we harvested B-ALL cells from the spleen, bone marrow, and meninges of leukemic mice and sorted them on the basis of their constitutive RFP expression levels to examine editing outcomes in variant-harboring cells **(Fig. 1i)**. To assess the correlation between reporter and endogenous gene editing activities, as well as determine whether we could identify leukemia cells expressing impactful mutations even if these are present at low frequencies within heterogeneous cell populations, we introduced a validated *Trp53*-targeting sgRNA in CBE reporter cells designed to install a C132Y (Cysteine to Tyrosine) mutation previously shown to impact p53 protein function^10^ (**Extended Data Fig. 3a-c**). As expected, we observed that edited cells harboring a *Trp53* mutant genotype outcompeted WT cells in both *in vitro* and *in vivo* competition assays (**Fig. 1j and Extended Data Fig. 3c**). Furthermore, loss of p53 function conferred resistance to treatment with Nutlin-3a and doxorubicin, which stabilize p53 or induce p53 activity, respectively (**Extended Data Fig. 3d**)^87,88^. Collectively, these proof-of-concept data demonstrate that our base editing *BCR-ABL+* B-ALL model could be used for high-throughput base editing studies to assess the impact of cancer-associated variants *in vivo*.

### Multiplexed *in vivo* base editing screens identify contextual and variant-specific phenotypes

A number of studies have shown that the effects produced by genetic perturbations can vary depending on the context^46–76^. To test the hypothesis that the impact of disease-associated genetic variants depends on one or more contextual variables, we separately introduced ABE and CBE sub-pools derived from MBESv2 libraries into B-ALL cells to perform parallel *in vitro* and *in vivo* screens (**Fig. 1f, 2a, see also below**). Library-transduced B-ALL cells harboring base editing activity reporters were electroporated with optimized ABE8e or CBE6 mRNAs and cultured for two days followed by purification of editing-competent GFP+ cells using fluorescence activated cell sorting. Five days later, library-transduced GFP+ cells (at ≥ 1,000X library representation) were cultured *in vitro* or transplanted into immunocompetent C57BL/6J recipient animals (**see Methods**). Next, we harvested the spleen, bone marrow, and meninges from mice at the end-stage of disease (Day 10) along with cells from time-matched *in vitro* screen counterparts followed by genomic DNA isolation, next-generation sequencing of integrated base editing sgRNA-sensor constructs, and computational analysis of count data to deconvolute differences in sgRNA/mutation behavior (log2-base fold changes, or LFCs, and Z-score calculations; see **Fig. 1f, 2a, and Methods**).

**Figure 2.**
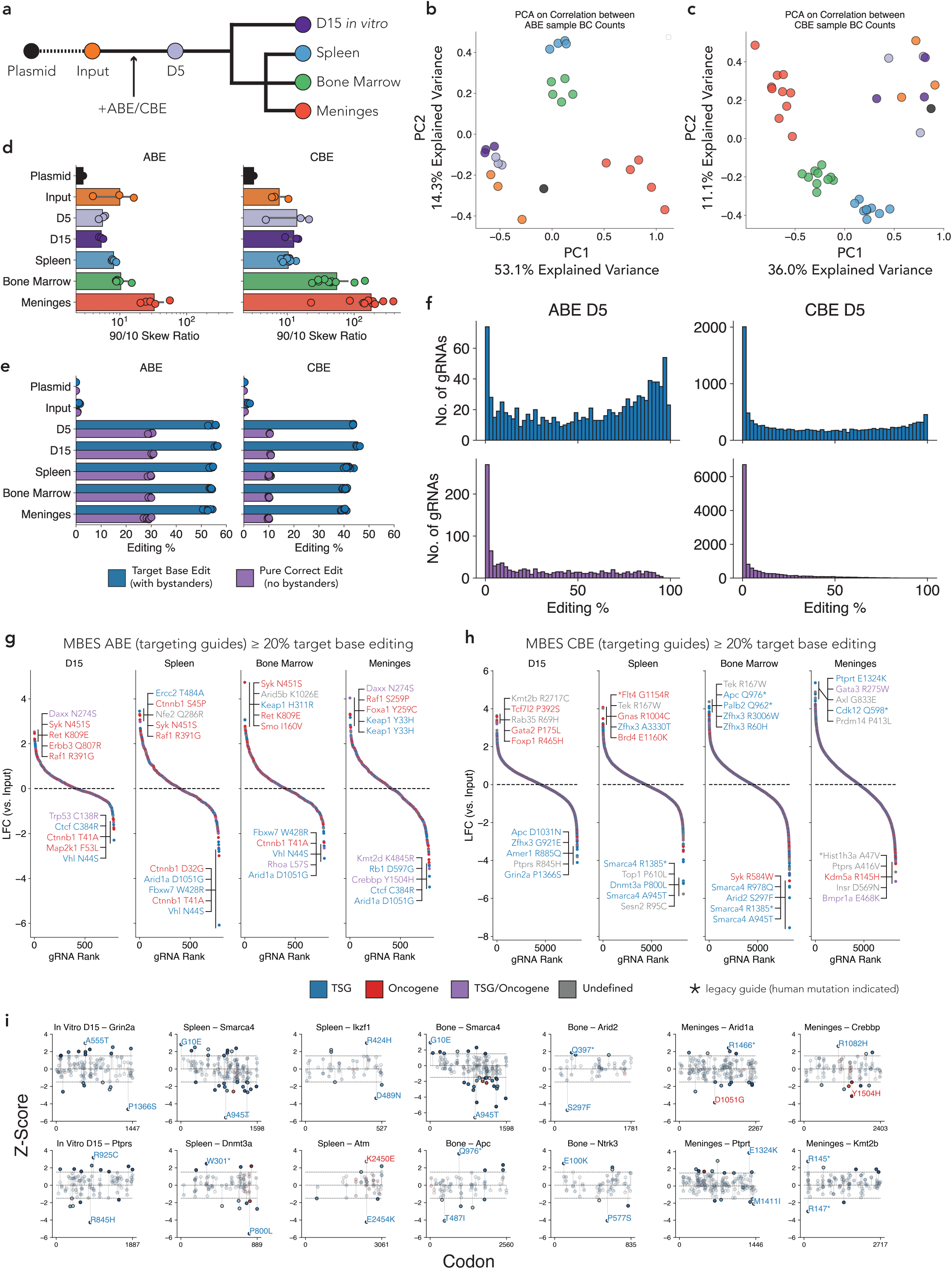
Multiplexed *in vivo* base editing screens identify contextual and variant-specific phenotypes. **a,** Schematic of samples generated from screening. **b,** Principal components analysis of the correlation matrix of barcode counts for replicates of indicated ABE samples. **c,** Same as **(b)**, but for CBE samples. **d,** The 90/10 skew ratio (counts of 90th percentile guide/counts of 10th percentile guide) for each indicated sample for ABE screen (left) and CBE screen (right). **e,** Average target base editing % (with bystander edits allowed, blue) and pure correct editing % (no bystander edits allowed, purple) at the sensor locus. **f,** Histogram of target base editing rate (top, blue) and pure correct editing rate (bottom, purple) at the sensor locus for the D5 samples. **g,** Waterfall plot of ABE gRNA enrichment in each sample type, with top 5 and bottom 5 guides labeled. Guides with fewer than 50 average control counts and target base editing <20% excluded. Points colored by COSMIC Cancer Gene Census annotation. **h,** Same as **(g),** but for CBE gRNAs. **i,** Lollipop plots of selected genes with the largest range of gRNA behaviors, showing gRNAs with ≥20% target base editing at the sensor locus, and labeling the top and bottom enriched and depleted gRNAs. Guides colored by their target base editing rate (ABE, red; CBE, blue), with gRNAs with a Z-score of ≤ −1.5 or ≥ 1.5 highlighted. See also Extended Figures 4-6.

To determine the extent to which genetic variants differentially influence the *in vivo* growth and systemic organotropism of cancer cells, we performed principal component analysis (PCA) of correlation scores among sgRNA read counts obtained from all sample groups (including *in vitro* screen replicates and individual mice). We observed consistent differences in sgRNA behavior between each biological context. For instance, while mutant leukemia cell populations growing *in vitro* were highly similar to each other, those growing in the spleen and bone marrow were more similar to each other than to any *in vitro* or meninges-derived leukemia cells (**Fig. 2b, 2c**). These analyses suggested that the physiological *in vivo* environment and tissue site are important contextual determinants of the behavior of specific genetic variants.

Although we and others have previously shown that the *BCR-ABL+* B-ALL model allows for robust representation of complex genetic libraries *in vivo*, we reasoned that some of these contextual differences may in principle arise as a consequence of *in vivo* genetic bottlenecking, in part due to the stochastic nature of cancer cell engraftment^89^. To examine this possibility, we assessed genetic bottlenecking by calculating the representation and frequency of ABE and CBE variants in each of these datasets. Skew ratios calculated using the distribution of 90th and 10th percentile sgRNAs (90/10) showed a relatively normal distribution of variants across tissues (**Fig. 2d**, **Extended Data Fig. 4a, 4b)**. As expected, the meninges exhibited moderately higher skew ratios compared to the other tissues examined (**Fig. 2d**). Higher skew ratios in this anatomical compartment could be influenced by limited leukemia cell occupation and lower engraftment rates relative to other tissue types like the spleen and bone marrow, leading to disproportional representation of some sgRNAs. To investigate this further, we quantified the total amount of sequencing reads assigned to each sgRNA and found that most reads obtained from different tissue compartments originated from diverse guides, with the exception of one CBE guide that encompassed up to 5% of the reads obtained from one meninges sample (**Extended Data Fig. 4c, 4d**). Notably, while some skewing exists in the meninges, the broad representation of sgRNAs in this context support a model by which diverse leukemia cell populations — and not just small numbers of metastatic cells — gain access to the meninges via osteoblastic breach of proximal skull tissue^83,84,90,91^. Taken together, our data suggest that this B-ALL model can be coupled with high-throughput base editing to explore how diverse endogenous genetic variants influence cancer cell behavior *in vitro* and in distinct anatomical *in vivo* contexts, including tissues where achieving and maintaining a high representation of complex genetic perturbations remains challenging for large-scale genetic screens (e.g. central nervous system; meninges). Below, we unpack these foundational datasets to illustrate the functional richness of gene- and mutation-specific phenotypes across several contexts.

### Dynamic calibration of *in vivo* base editing screens identifies a functionally diverse mutational compendium of cancer drivers and dependencies

A key advantage of our *in vivo* base editing screening framework is the inclusion of variant-specific synthetic sensors in our libraries (**Fig. 1e, Extended Data Fig. 1g**). In contrast to standard CRISPR and base editing screens that rely on sgRNA count data to indirectly assess enrichment and depletion of putative endogenous alleles^24,32,33,35,40,92^, our sensors allow for quantitative and dynamic (re)calibration of multiplexed base or prime editing screening data to uncover guide RNAs and variants whose phenotypic output or severity may vary depending on the degree and nature of diverse types of editing events. In doing so, these sensors allow us to simultaneously measure the fitness effects, mutational patterns, and overall editing efficiency and precision of diverse sgRNAs and variants in high-throughput across various experimental contexts (**Fig. 1e and Extended Data Fig. 1g, 1h**).

We measured base editing efficiency using two metrics: ‘target’ base editing and ‘perfect’ base editing (**Fig. 2e, 2f**). Target base editing quantification takes into account bystander mutations within the optimal sgRNA editing window, while perfect base editing exclusively quantifies the activity at intended nucleotides. Depending on the gene and mutation being targeted, some instances of target base editing are equal to perfect base editing frequencies. On average, sensor editing was detected in >40% of sequencing reads, with higher editing being observed in the ABE screen (**Fig. 2e**). This finding is consistent with published and anecdotal data showing that ABE8e is a more efficient genome editor compared to CBE6^81,82^. Moreover, target base editing was highest within the +5 to +7 region of the optimal editing window, consistent with previous studies (**Extended Data Fig. 5a**)^81,82^. Importantly, editing was highly correlated and similarly distributed across all experimental conditions (**Extended Data Fig. 5b-d**), highlighting platform and screening robustness.

As stated above, our sensors allow rigorous filtering of screening data based on a dynamic threshold of editing efficiency and precision reported as percentages of target and perfect base edits (**Extended Data Fig. 1h**). Our previous work showed that sensor-based target editing efficiencies of >20% were sufficient to quantitatively assess variant-dependent fitness effects (based on corresponding sgRNA/mutation LFCs) (**Figure 2f**), though we want to stress that this is a dynamic functional threshold akin to an analog lever of editing^10,11^ (**Extended Data Fig. 1h**). Using this threshold, we identified 802/1,034 ABE and 8,449/12,806 CBE mutation-specific sgRNAs targeting a broad spectrum of genes known or suspected to act as oncogenic drivers, dependencies, or modulators of cancer phenotypes and therapeutic outcomes in diverse types of human leukemias and hematopoietic malignancies, as well as in a wide spectrum of epithelial cancers (**Fig. 2g, 2h**). Many of these genes and mutations are the targets of experimental or clinical therapies (including FDA approved drugs), suggesting that this approach can be used to nominate new drug targets.

Our mutational compendium is functionally diverse and includes both positively- and negatively-selected variants distributed across well-defined gene categories, such as kinases, transcription factors, chromatin regulators, RNA binding proteins and splicing factors, DNA polymerases, translation initiation factors, immune receptors, scaffolding proteins, G-protein coupled receptors, guanine nucleotide exchange factors, GTPase-activating proteins, and ubiquitin enzymes, among many others (**Fig. 2, 3, Extended Data Fig. 6**). For instance, we observed positive-selection for mutant alleles in many kinases (*Syk, Ret, Erbb2, Erbb3, Egfr, Pik3r2, Pik3r3, Pik3cg, Mtor, Rictor, Rptor, Raf1, Araf, Braf, Map2k1/Mek1, Map3k1/Mekk1, Akt1, Akt3, Ntrk2, Ntrk3, Fgfr2, Fgfr3, Jak1, Cdk4, Axl, Kit, Yes1, Kdr, Lck, Lyn, Flt1, Flt3, Flt4, Acvr1, Lats1, Rps6ka4, Atm, Chek2, Pim1*), phosphatases (*Ptprt, Ptprs, Ptpn11*), non-enzymatic kinase regulators (*Tsc1, Tsc2, Card11, Socs1*), growth factors, cytokines, their receptors, and signaling regulators (*Fgf2, Fgf3, Irs1, Irs2, Csf3r, Cd79a, Epha5, Epha7, Ephb1, Myd88*), transcription factors, co-activators, and key regulators (*Pax5, Myc, Mga, Fubp1, Ncoa3, Cebpa, Tp53, Runx1, Smad4, Rara, Gata2, Gata3, Etv6, Ikzf1, Stat3, Irf8, Sox9, Tbx3, Erg, Ar, Zfhx3, Nkx2-1, Nfe2l2, Yap1, Gli1, Pgr, Id3, Tcf7l2*), G-protein coupled receptors, their regulators, and other transmembrane receptors (*Smo, Ptch1, Notch1, Notch3, Nf2, Gnas*), RNA binding and processing proteins (*Rbm10, U2af1, Ago2, Dicer1, Msi2, Spen, Csde1, Dis3*), translation factors (*Eif1ad19, Eif4a2*), GTPases and their regulators (*Kras, Rras1, Rras2, Rheb, Rab35, Nf1*), and metabolic enzymes (*Idh1, Idh2, Fh1, Inha*). We also found significant enrichment of mutations in functional gene groups involved in key cellular and molecular processes, including enzymatic and non-enzymatic chromatin regulators (*Kmt2a, Kmt2b, Kmt2d, Kdm6a, Men1, Atrx, Daxx, Ep300, Crebbp, Smarca4, Arid5b, Carm1, Brd4, Nsd1, Bcor*), DNA modification enzymes (*Dnmt3a, Tet2*), regulators of genome topology and compaction (*Ctcf, Stag2*), DNA damage, repair, and maintenance sensors and effectors (*Rad50, Ercc3, Palb2, Rtel1, Atm, Bap1, Atrx, Daxx, Pole*), and ubiquitin activating, conjugating enzymes, and ligating enzymes (*Keap1, Cul3, Traf7, Fbxw7, Vhl*), as well as several other multi-functional genes and proteins involved in a diversity of important pathways. These results demonstrate that our multiplexed base editing platform can represent highly complex libraries of genetic perturbations *in vivo* and allows robust, functional interrogation of a broad spectrum of mechanistically diverse genetic variants in genes involved in diverse cellular and molecular processes.

**Figure 3.**
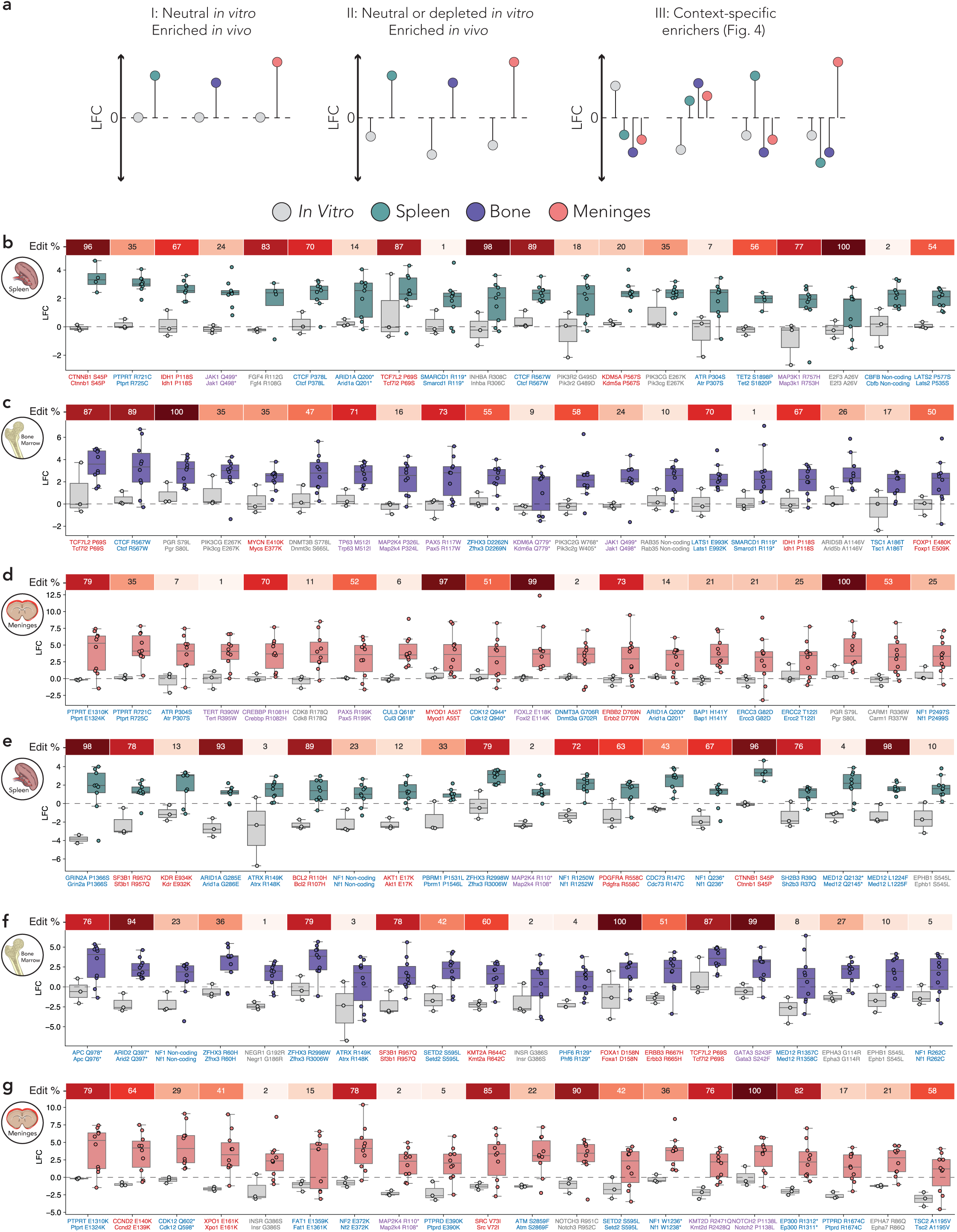
Base editing screens uncover functional and contextual phenotypic divergence of genetic variants. **a,** Schematic of the different classes of gRNA behavior that we identified in our screen. **b,** Spleen vs. *in vitro* gRNA enrichment, showing gRNAs with −0.25 ≤ LFC ≤ 0.25 *in vitro*, and enriched in the spleen, ranked in order of difference between spleen and *in vitro* LFC. **c,** Same as **(b),** but for bone marrow vs. *in vitro* comparison. **d,** Same as **(b)**, but for meninges vs. *in vitro* comparison. **e,** Spleen vs. *in vitro* gRNA enrichment, showing gRNAs with LFC<0 *in vitro*, and enriched in the spleen, ranked in order of difference between spleen and *in vitro* LFC. **f,** Same as (e), but for bone marrow vs. *in vitro* comparison. **g,** Same as (e), but for meninges vs. *in vitro* comparison. See also Extended Figures 6-7.

### Divergent phenotypes produced by gene-specific variants and genetic knockouts

It is important to highlight that positively-selected mutations likely encompass a diverse mix of gain-of-function (GOF) and loss-of-function (LOF) alleles, and may also include separation-of-function, change-of-function (neomorphic), and dominant negative mutations whose functional impact remains unknown. Other than LOF, these types of mutant alleles may be fundamentally different to those obtained using traditional gene disruption technologies like RNA interference (RNAi) and CRISPR-Cas9 nuclease approaches, where positively-selected events almost always map to genes that suppress cellular fitness (e.g. tumor suppressor genes). To formally test this hypothesis, we conducted a parallel complementary CRISPR-Cas9 nuclease screen using the ABE MBESv2 library, essentially as described **in Figure 1**. Notably, sgRNA LFC values obtained from the CRISPR-Cas9 nuclease screen were poorly correlated with those obtained from the ABE screen across all conditions (**Extended Data Fig. 7a**). This was not a technical failure of CRISPR-Cas9 nuclease screens, as we observed similar correlation patterns among samples in the Cas9 nuclease screen, with the strongest correlation observed between the spleen and bone marrow samples (**Extended Data Fig. 7b**). These results suggest that CRISPR-Cas9 nuclease screens can accurately model genetic LOF phenotypes but are largely unable to recapitulate specific genetic events that may confer cellular and molecular GOF, change-of-function, or separation-of-function phenotypes, or that alter protein conformation or binding capabilities. For instance, Cas9-mediated disruption of a proto-oncogene would likely produce fundamentally different effects to base or prime editing-mediated engineering of an oncogenic GOF missense variant. Consistent with this hypothesis, we found that sgRNAs that exhibited the most divergent behavior across ABE and Cas9 screens targeted a functionally diverse set of genes that encode for kinases (e.g. *Raf1, Syk*), transcription factors (e.g. *Etv6, Sox9, Foxo1*), gene expression and chromatin regulators (e.g. *Daxx, Dnmt3a*), and others (**Extended Data Fig. 7c**). Together, these results suggest that the functional impact of many specific endogenous genetic variants cannot be assessed with standard CRISPR-Cas9 nuclease approaches.

### Base editing screens uncover functional and contextual phenotypic divergence of genetic variants

As discussed above, a key aspect of our approach that distinguishes it from prior work in the field is that we interrogate the phenotypes produced by sgRNAs that introduce specific gene *mutations* instead of gene disruption or silencing (e.g. using Cas9 nucleases or CRISPR interference). Doing so allows us to probe the functional impact of individual mutations affecting the same gene(s) to determine whether different mutations within a gene may produce divergent phenotypes, and whether these effects could vary depending on the context in which the mutations arise or act. To test this concept, we analyzed our mutational compendium to identify and distinguish between three categories of genes and mutations (**Fig. 3a**): (1) those that predominantly exhibit *opposing* phenotypes depending on the location within the gene and the type of mutation, (2) those that predominantly exhibit *neutral* phenotypes in one context but strong selection in more than one context, and (3) those that predominantly exhibit *context-specific* phenotypes. To identify and distinguish between these three categories, we performed a series of comparative data analyses between one or more contexts using median LFC and Z-score values for gene-specific sgRNAs designed to introduce specific mutations in different regions of the gene (**see Methods**). The results of these analyses are shown in **Figure 2i**, **Figures 3-4**, and discussed below.

**Figure 4.**
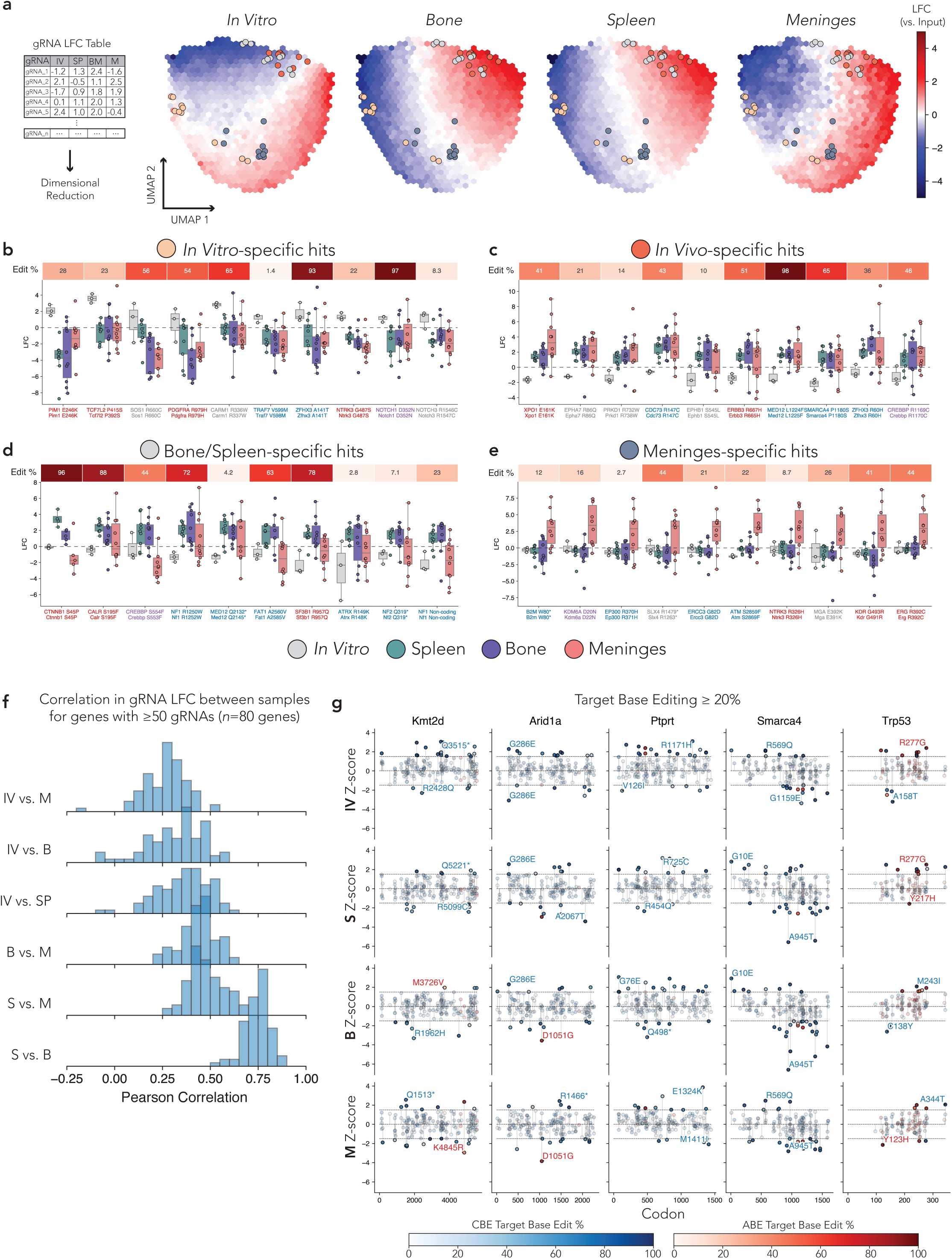
Comparative *in vivo* base editing screens nominate tissue-specific and organotropic genetic variants. **a,** The LFC of each gRNA in each tissue (4-dimensional vector) was dimensionally reduced for visualization on a UMAP plot. Each plot is a heatmap showing the LFC of gRNAs in each indicated tissue type in UMAP space. Points, and their colors, correspond with the four sets of gRNAs selected in figures **(b-e)**. **b,** Boxplot of the LFC of gRNAs in each replicate for selected *in vitro*-specific hits. Target base editing rate at the sensor locus shown at top of plot. Guides labeled with targeted human mutation (top) and corresponding mouse mutation (bottom), and colored by COSMIC Cancer Gene Census annotation (red, oncogene; blue, TSG; purple, TSG/oncogene; grey, unannotated). **c,** Same as **(b)**, but for selected *in vivo-*specific hits. **d,** Same as **(b)**, but for selected bone/spleen*-*specific hits. **e,** Same as **(b)**, but for selected meninges*-*specific hits. **f,** Histogram of the Pearson correlation for gRNA LFC in a given gene between samples, for genes with ≥50 gRNAs (*n*=80 genes). IV, *in vitro*; S, spleen; B, bone marrow; M, meninges. **g,** Lollipop plots of the top 5 represented genes in the screen, showing gRNAs with ≥20% target base editing at the sensor locus, and labeling the top and bottom enriched and depleted gRNAs. Guides colored by their target base editing rate (ABE, red; CBE, blue), with gRNAs with a Z-score of ≤-1.5 or ≥1.5 highlighted. See also Extended Figures 6-7.

To identify genes and mutations that predominantly exhibit *opposing* phenotypes depending on the location and type of mutation, we examined all gene-specific sgRNAs (mutations) across all contexts (*in vitro*, spleen, bone marrow, meninges). This analysis identified several genes and mutations, including *Grin2a, Smarca4, Ikzf1, Arid2, Arid1a, Crebbp, Ptprs, Dnmt3a, Atm, Apc, Ntrk3, Ptprt,* and *Kmt2b*, among others (**Fig. 2i**). Intriguingly, these analyses also identified putative functionally opposing nonsense mutations in *Kmt2b* that were separated by only two residues: *Kmt2b.R145\** and *Kmt2b.R147\** (* denotes a premature termination codon, or PTC) (**Fig. 2i, bottom right panel**). These results suggested that the R145* nonsense mutation was positively selected (not detrimental to cells) while the R147* nonsense mutation was negatively selected (detrimental to cells). As this was unexpected, we decided to examine the KMT2B protein sequences flanking each of these residues to determine whether there was anything special about them. The only notable difference we found is that the R147 (Arg^147^) residue is preceded by a glycine (Gly) at position 146 (Leu^141^–Arg^142^–Ser^143^–Gln^144^–Arg^145^–**Gly**^146^–PTC), while R145 (Arg^145^) is preceded by a glutamine (Gln) at position 144 (Leu^141^–Arg^142^–Ser^143^–**Gln**^144^–PTC). Intriguingly, recent work from Jagannathan and colleagues^93^ has shown that the Gly-PTC mRNA context is a highly efficient trigger of nonsense mediated decay (NMD). These results suggest the testable hypothesis that the *Kmt2b.R147\** nonsense mutation may be detrimental to cells due to being a highly efficient target for NMD, while the *Kmt2b.R145*\* nonsense mutation may escape NMD and be tolerated by cells. More broadly, these results highlight the unique resolution and sensitivity of our approach to potentially distinguish between functionally opposing mutations affecting sites that are right next to each other, which would otherwise be classified as functionally equivalent merely based on physical proximity.

To identify genes and mutations that may predominantly exhibit *neutral* phenotypes in one context but strong selection in more than one context, we first looked for sgRNAs that produced little-to-no fitness differences *in vitro* (defined as having a ‘neutral’ median LFC between −0.25 and +0.25) but that exhibited strong positive-selection in at least one *in vivo* context (e.g. neutral behavior *in vitro* but strong enrichment in the spleen and bone marrow) (**Fig. 3a**). These analyses identified several genes and mutations that exhibited this behavior in each of the three *in vivo* contexts (**Fig. 3**). For instance, *in vitro* vs spleen analyses identified mutations in *Ctnnb1, Ptprt, Idh1, Jak1, Fgf4, Ctcf, Arid1a, Tcf7l2, Smarcd1, Inhba*, and others (**Fig. 3b**), *in vitro* vs bone marrow identified mutations in *Tcf7l2, Ctcf, Pgr, Pik3cg, Mycn, Dnmt3b, Trp63, Map2k4, Pax5, Zfhx3*, and others (**Fig. 3c**), and *in vitro* vs meninges identified mutations in *Ptprt, Atr, Tert, Crebbp, Cdk8, Pax5, Cul3, Myod1, Cdk12,* and *Foxl2,* among others (**Fig. 3d**). We also extended our search to look for sgRNAs that produced strong opposing fitness differences when comparing *in vitro* effects to any *in vivo* contexts (e.g. negative selection *in vitro* but strong positive selection in the spleen). These analyses identified mutations in *Grin2a, Sf3b1, Kdr, Arid1a, Atrx, Bcl2, Nf1, Akt1, Pbrm1, Zfhx3,* and others in the spleen (**Fig. 3e**), *Apc, Arid2, Nf1, Zfhx3, Negr1, Atrx, Sf3b1, Setd2, Kmt2a, Phf6,* and others in the bone marrow (**Fig. 3f**), and *Ptprt, Ccnd2, Cdk12, Xpo1, Insr, Fat1, Nf2, Map2k4/Mkk4, Ptprd, Src,* and others in the meninges (**Fig. 3g**). Each of these genes and their mutations have been linked with various types of human cancers, including hematopoietic malignancies (*Sf3b1, Arid1a, Bcl2, Setd2, Kmt2a, Phf6, Xpo1*) and many other pediatric and adult cancer subtypes. Together, these results demonstrate the potential of our platform to identify functionally impactful mutations that exhibit broad malignant potential across diverse disease contexts.

### Comparative *in vivo* base editing screens nominate tissue-specific and organotropic genetic variants

The functional and phenotypic richness of the above results prompted us to explore the possibility that the cellular environment is a major contextual modulator of the behavior and functional selection of gene-specific mutations. To visualize the landscape of differences in sgRNA enrichment and depletion in the different tissues and contexts, we performed Uniform Manifold Approximation and Projection (UMAP) on the matrix of LFCs of sgRNAs by context (*in vitro*, spleen, bone marrow, meninges) (**Fig. 4a, see Methods**). This analysis and visualization showed discernible patterns of co-enrichment and co-depletion for groups of sgRNAs, but also clusters of sgRNAs that exhibited context-specific differences. We also noticed that a subset of sgRNAs may exhibit tissue- and context-specific enrichment patterns (**Fig. 4b-d**), consistent with the hypothesis stated above. We reasoned that, if this is true, we could potentially identify genes and mutations that *only* exhibit *context-specific* phenotypes (e.g. those that exhibit strong positive selection only in one context).

To test this hypothesis, we performed systematic ‘gene-by-variant-by-context’ analyses to determine whether any variants exhibit detectable contextual phenotypes (**Fig. 4**, **5**, **Extended Data Fig. 8**). First, we focused our analysis on *in vitro* vs *in vivo* differences. Because we examined the fitness effects of variants across three tissues/organs, we first compared *in vitro* with pooled *in vivo* samples. We first looked for sgRNAs that did not exhibit positive selection *in vitro* (median LFC < 0) but did so *in vivo* (median LFC > 1 for all tissues). This analysis identified several genes and mutations that exhibit positive selection *in vivo*, including *Xpo1, Epha7, Prkd1, Cdc73, Ephb1, Erbb3, Med12, Smarca4, Zfhx3,* and *Crebbp* (**Fig. 4c**). Notably, the most differentially enriched genetic variant, *Xpo1.E161K*, was recently found to be an oncogenic driver in human B-cell malignancies that is altered through a recurrent change-of-function mutation in the *XPO1* nuclear export receptor^94^. Based on the finding that cancer cells driven by the XPO1.E161K mutation are hyper-sensitive to XPO1 inhibitors, FDA-approved small molecule inhibitors of XPO1 (e.g. selinexor) are currently being investigated in several clinical trials^95–101^. Molecularly, XPO1 has also been linked to micronuclear rupture and chromosomal instability, suggesting broader cancer relevance^102^.

**Figure 5.**
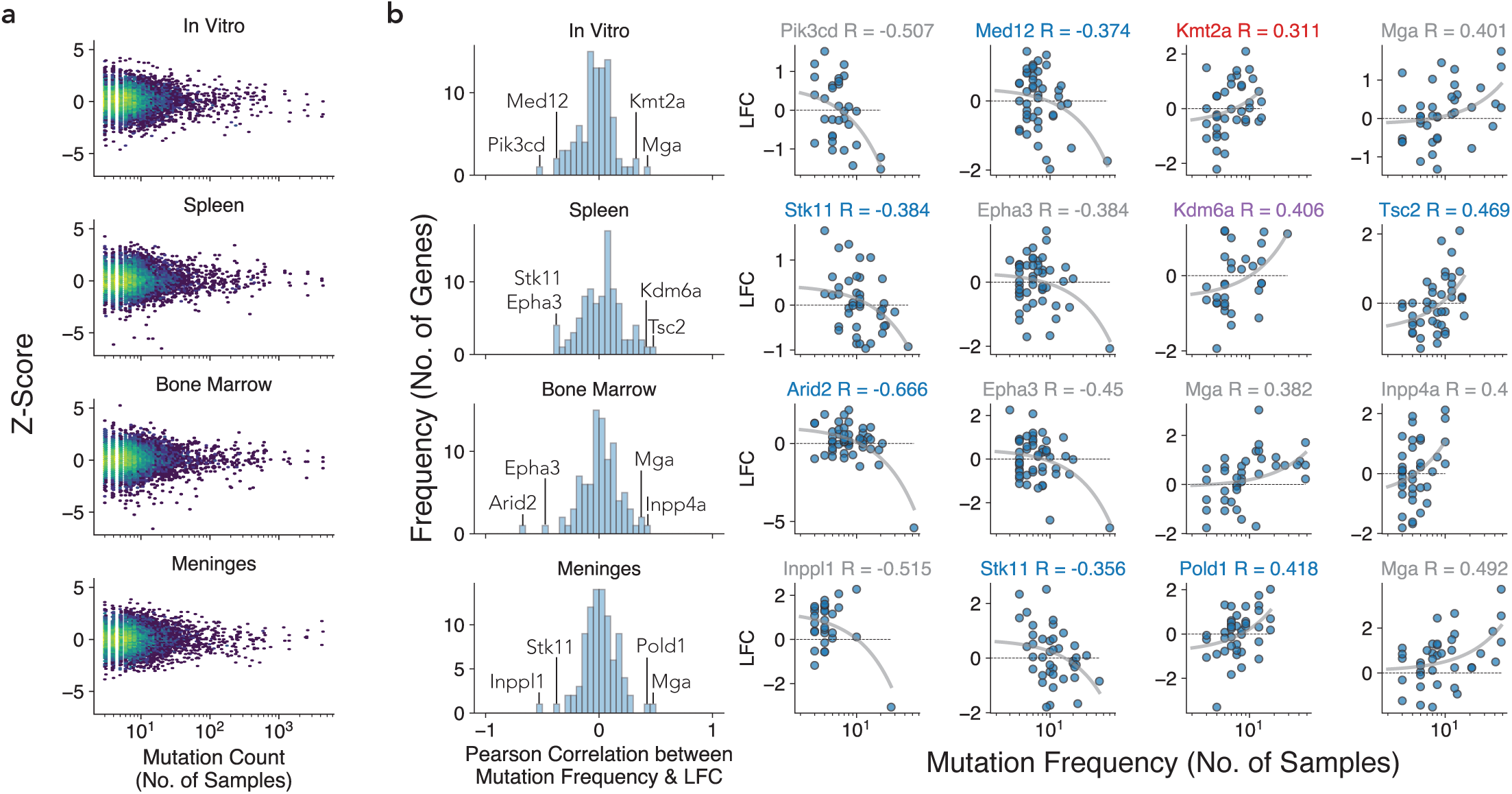
Mutation frequencies of genetic variants are not always correlated with functional impact. **a,** Scatterplot of the mutational frequency (number of samples where mutation is observed) and LFC of the corresponding gRNA designed to engineer that mutation for each tissue/context. **b,** Histogram of the Pearson correlation between mutational frequency and LFC on a *per gene* basis, separated by tissues. Analysis restricted to gRNAs with target base edit ≥ 20% and genes with at least 30 gRNAs above this editing threshold (n = 92 genes). The top and bottom 2 genes, as ranked by Pearson correlation with mutational frequency, are plotted at right, with linear regression line shown in grey.

We then performed reciprocal analyses to identify sgRNAs that exhibited positive selection *in vitro* (median LFC > 1) but did not exhibit positive selection *in vivo* (median LFC < 0 for all tissues). This analysis identified several genes and mutations, including *Pim1, Tcf7l2, Carm1, Notch1,* and *Notch3*, among others (**Fig. 4b**). Of note, activating *Notch1* and *Notch3* mutations that drive aberrant oncogenic signaling are commonly observed in T-cell acute leukemia (T-ALL) patients, yet these same genetic lesions have been shown to be incompatible with fitness and survival of B-ALL cells unless they acquire additional mutations that promote cellular plasticity and transition to a T-cell lineage (e.g. *Phf6* mutations)^103,104^. More broadly, the observation that *in vitro* perturbations exhibit divergent behavior to corresponding *in vivo* phenotypes is consistent with our prior work (**Fig. 3b, 3c**)^50,56,57^. These bidirectional analyses suggest that our approach can identify specific genes and mutations that exhibit contextual phenotypes, including those that influence cell state and lineage transitions.

Having observed *in vitro* vs *in vivo* functional differences, we then wondered whether specific genes and mutations may exhibit organ-specific positive selection patterns. The spleen and bone marrow are critical organs for lymphoid cell development and immune function, and are common sites for leukemia cell infiltration^83^. In line with their functional and pathological similarities, we observed similar genetic variant correlations between these tissue types, suggesting that these cellular compartments are similar enough to not drive significant divergence in the behavior of the genetic interactions modeled (**Fig. 4d, Extended Data Fig. 8**). Thus, because spleen and bone marrow samples were very highly correlated, we decided to group them together for the analyses described below (‘SP-BM’). On the other hand, because B-ALL frequently disseminates to the central nervous system (CNS), particularly the meninges^84^, we reasoned that this type of analysis may uncover putative genetic factors that uniquely contribute to meningeal infiltration in B-ALL.

To investigate the possibility that certain genes and mutations may exhibit organ-specific positive selection patterns, we performed systematic pairwise analyses looking for sgRNAs that exhibited positive selection *predominantly in specific tissues* (median LFC > 1 for one tissue) but did not exhibit positive selection *in any other context* (median LFC < 0 for all other contexts) (**Fig. 4**). Comparative analyses of SP-BM samples with all other contexts, as well as meninges samples with every other context, identified several genes and mutations that showed positive selection predominantly in the spleen and bone marrow (**Fig. 4d**), or in the meninges (**Fig. 4e**). For instance, we found that certain mutations in *Ctnnb1, Crebbp, Fat1*, *Sf3b1*, and *Nf1* predominantly exhibit positive selection in the spleen and bone marrow, while other mutations in *B2m, Kdm6a, Ep300, Slx4, Ercc3, Atm, Ntrk3, Mga, Kdr,* and *Erg* were found to be particularly enriched in meningeal tissue. Intriguingly, some of these genes have been established or hypothesized to exhibit mutual functional and contextual interactions based on prior work. For instance, *Slx4* mutations have been associated with brain metastasis^105^, and SLX4 has been shown to interact with MGA, ERCC1 (another ERCC family member), and CDC73 (encoded by *Cdc73*, whose mutant alleles also exhibited positive selection *in vivo*)^106^ (**Fig. 4c**), while interactions between the proteins encoded by *Ep300* and *Kdm6a* are involved in enhancer regulation^107^.

To more broadly examine the functional effects of gene-by-gene and variant-by-variant relationships across contexts, we identified genes that were targeted by the highest number of sgRNAs (and thus mutagenized much deeper than all other genes). We identified 80 genes that had a minimum of 50 sgRNAs designed to install a spectrum of genetic variants across the gene body (**Fig. 4f**). To assess the potential degree of functional heterogeneity on a gene-by-variant-by-context level, we looked at whether the behavior of individual variants fluctuated within and across each of the contexts. In line with global analyses, spleen and bone marrow samples were highly correlated to each other in comparison to the meninges, while *in vitro* samples exhibited low correlation to all tissue types (**Fig. 4f**). We then focused our analysis on a subset of genes that were closer to saturation across all screens: *Kmt2d, Arid1a, Ptprt, Smarca4,* and *Trp53*. Consistent with the results obtained from the analysis of genes and mutations that exhibit opposing phenotypes depending on the location and type of mutation (**Fig. 2i**), we observed notable differences in the pattern of enriched and depleted variants on a per-gene basis across all tissue types (**Fig. 4g**). These data suggest the existence of gene-by-variant-by-context interactions that could play a functional role in modulating tissue-specific cancer cell survival or the organotropic behavior of cancer cells, which should be the subject of future investigation.

### The observed mutation frequency of a genetic variant is not always correlated with functional impact

The mutation frequency of a variant is a function of the (1) probability of the variant emerging by chance, (2) the pathogenicity of the variant, (3) the immunogenicity of a variant, and (4) the context-specific behavior of a variant. To examine this in more detail, we determined the extent to which the observed frequency of a specific mutation correlates with the cellular fitness impact observed in our screens. We reasoned that, if variant pathogenicity is the major determinant of fitness effects, we would observe a clear correlation between a variant’s mutation frequency and variant-specific sgRNA enrichment or depletion. In contrast, we observed that the LFC of variant-specific sgRNAs does not generally correlate with the observed mutation frequencies (**Fig. 5a**). Intriguingly, for a subset of genes and mutations (e.g *Kmt2a, Mga, Kdm6a, Tsc2, Inpp4a,* and *Pold1*), we observed tissue-specific correlation patterns between the mutational frequency and degree of variant enrichment, suggesting putative gene-by-variant-by-context interactions (**Fig. 5b**). Taken together, these results illustrate the potential of our platform for deep functional and mechanistic analyses of endogenous genetic variants to disentangle complex biological phenomena across multiple physiological contexts.

## Discussion

The advent of targeted and whole genome sequencing technologies has made it feasible to routinely profile the genetic material of both healthy and sick individuals. Efforts from large-scale consortia like UK Biobank, 1000 Genomes Project, Pan-Cancer Analysis of Whole Genomes (PCAWG), Consortium of the International Cancer Genome Consortium (ICGC), The Cancer Genome Atlas (TCGA), and others have uncovered a vast array of genetic variants associated with ancestry and population-level human variation, but also thousands of alleles linked to disease susceptibility, development, and treatment responses, particularly in cancer^1–6^. While this information holds tremendous promise for improving clinical diagnoses and tailoring therapies based on patient genotype, the precise role and functional significance of most genetic variants remain unclear. Advanced genome editing technologies like base and prime editing offer the unique capability to systematically engineer and interrogate these genetic variants within their native genomic context. We and others have recently developed and applied scalable, sensor-based methods that allow for concomitant engineering and quantitative assessment of the phenotypic impact of most types of genetic variants^10–12^. However, high-throughput genome editing approaches have largely focused on investigating the impact of endogenous genetic variants on *in vitro* cellular phenotypes, thereby ignoring contextual and physiological *in vivo* pressures that may vary depending on the tissue or microenvironment. This is important, as several studies from our group and others have shown that the impact of a perturbation can be profoundly shaped by environmental and physiological contexts^46–63^. Moreover, while high-throughput gene essentially profiling efforts using RNAi^108,109^, CRISPR-Cas9 (refs. 110–112), CRISPRi^113^, CRISPRa^113^, ORF libraries^114^, and insertional mutagenesis^115^ have defined catalogs of disease-specific essential genes, these methods are largely unable to interrogate and distinguish the effects of specific types of mutant alleles that are observed in humans.

To address this challenge, we developed and applied new computational methods to design next-generation sgRNA libraries that accurately engineer evolutionarily conserved mutations (Dong et al., *in revision*) and screened these libraries *in vivo* by coupling hit-and-run base editing with immunocompetent mouse models capable of representing complex sgRNA libraries. By performing parallel *in vitro* and *in vivo* screens with a library of 13,840 sgRNAs to engineer 7,783 human cancer-associated mutations mapping to 489 endogenous protein-coding genes in mice, we generated a rich compendium of putative functional interactions between diverse categories of genes, mutations, and physiological contexts. We used this mutational compendium to (1) validate oncogenic and tumor suppressor gene mutations associated with diverse types of human cancers, (2) identify putative functionally-distinct mutations mapping to the same gene/protein predicted to exhibit different biochemical and biophysical activities, (3) uncover contextual and variant-specific phenotypes that may be shaped by the type of mutation, environment, and tissue type, and (4) nominate mutations that may influence cellular organotropism and meningeal infiltration *in vivo*. Many of the genes and mutations we identified are known or predicted to be druggable (OncoKB)^116,117^; indeed, some of these are currently being investigated in clinical trials to treat diverse types of human cancers^15,95,118–121^, while others are the targets of FDA-approved drugs^122–126^. Future work integrating this type of mutation-specific functional genetics framework with chemotherapies or targeted therapies could help identify modulators of therapeutic response.

Importantly, we also show that many mutations and their *in vivo* effects fail to be detected with standard CRISPR-Cas9 nuclease gene disruption approaches and, in some cases, produce discordant functional phenotypes. Indeed, while scalable gene disruption approaches are particularly effective at identifying genes that promote or suppress cellular fitness, they are unable to detect positive selection of mutationally-activated oncogenes and other types of drivers and dependencies. In contrast, we show that impactful oncogenic mutations that drive or otherwise modulate cellular fitness in different physiological contexts can be readily identified using our scalable multiplexed base editing approach. In practice, this converts what would otherwise be a ‘down assay’ with limited signal-to-noise ratios into an ‘up assay’ that may exhibit a wide dynamic range^127,128^ across challenging contexts, such as *in vivo* and within specific tissue microenvironments.

We envision this versatile platform and the generalizable framework described here could be readily deployed to investigate how genetic variation impacts *in vivo* phenotypes associated with cancer and other genetic diseases, as well as identify new potential therapeutic strategies to improve patient outcomes. More broadly, the modular, scalable, and multiplexed nature of our approach could be extended to interrogate more complex genetic interactions, such as gene-by-gene, gene-by-variant, and variant-by-variant relationships, dynamic interactions between each of these and the environment^129,130^, and higher-order epistatic relationships using multiplexed CRISPR effectors like AsCas12a^131^.

## Supporting information

Supplementary Protocol 1

Supplementary Protocol 1

## Extended Data Figure Legends

**Extended Data Figure 1.**
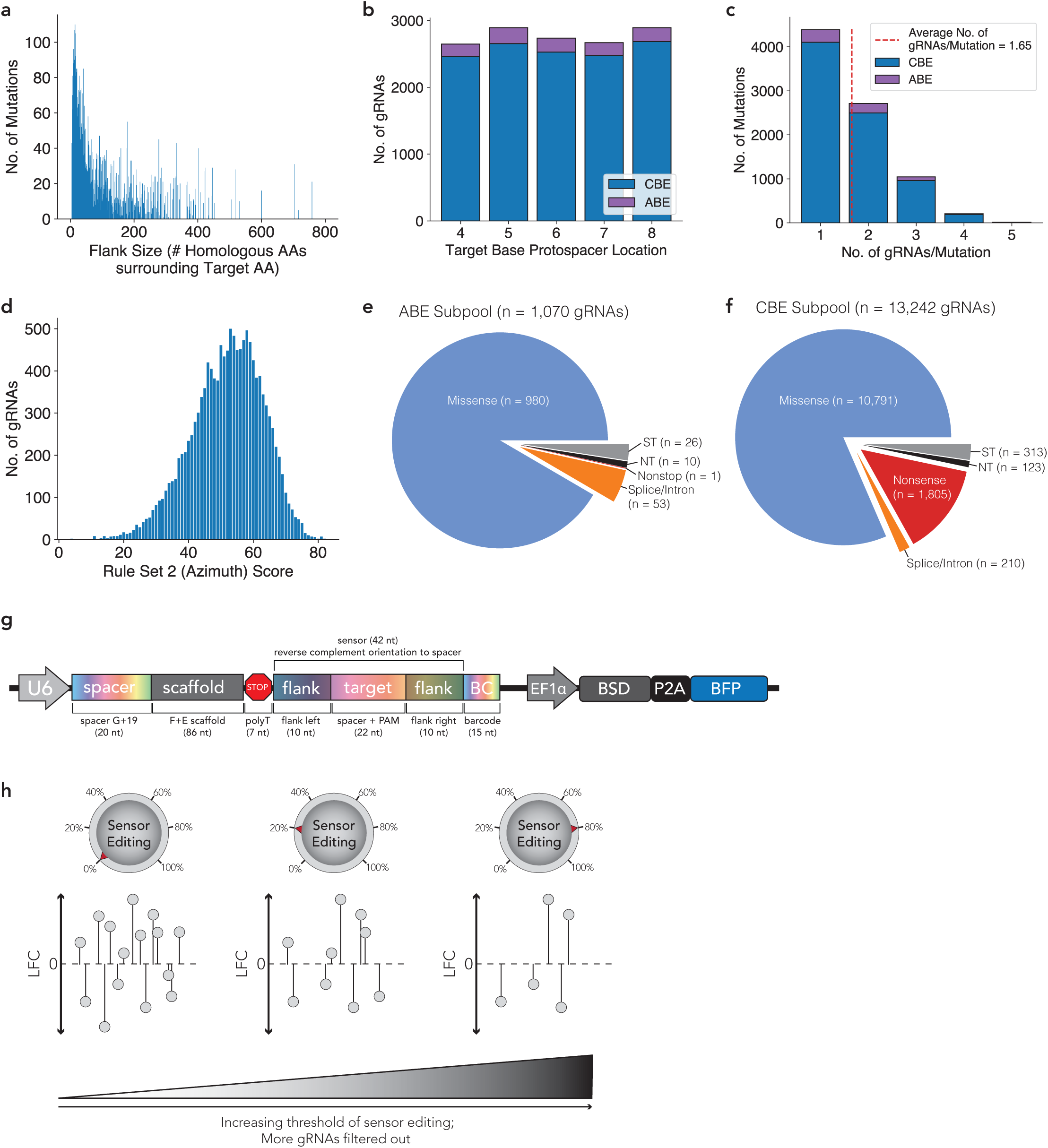
Summary description & analysis of MBESv2 library. **a,** Histogram of the “flank size,” the number of homologous amino acids between the human and mouse protein on either side of the target amino acid being mutated, for missense and nonsense mutations in the MBESv2 library (Dong et al., *in revision*). **b,** Histogram of the target base location within the protospacer for MBESv2 gRNAs. **c,** Histogram of the number of gRNAs per mutation. **d,** Histogram of the Rule Set 2 (Azimuth) on-target score for gRNAs included in the library. **e,** Pie chart of the ABE Subpool of MBESv2. NT, non-targeting guide; ST, safe-targeting guide. **f,** Pie chart of the CBE Subpool of MBESv2. NT, non-targeting guide; ST, safe-targeting guide. **g,** Detailed schematic of the sensor construct. BSD, blasticidin S resistance cassette; P2A, peptide 2ART; BFP, blue fluorescent protein. **h,** Sensor-based measurements provide a functionality akin to an analog lever of editing that can be used to establish dynamic editing thresholds.

**Extended Data Figure 2.**
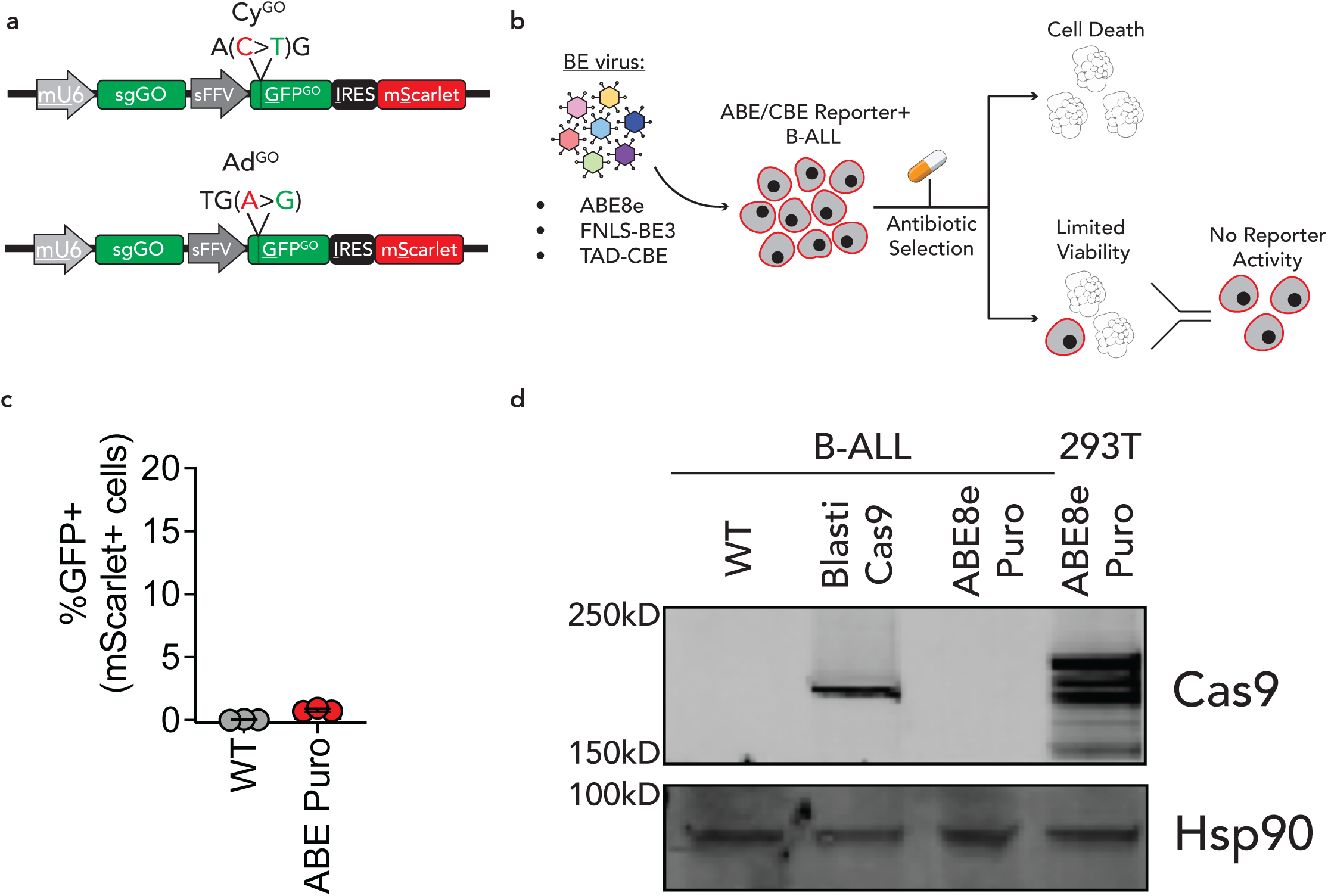
Attempts to stably express base editors in B-ALL cells. **a,** Schematic of GFP^ON^ (GO) base editing reporters utilized to quantify editing activity in B-ALL cells (adapted from Katti et al. 2020 *Nucleic Acids Research*; see ref. 85). Base editing enables production of GFP mRNA transcript by *de novo* creation of start codons (Cy^GO^) or removal of premature stop codons (Ad^GO^). **b,** Schematic of experimental strategy for stable expression of base editors in B-ALL cells. Cells either did not recover from antibiotic selection, or had low viability with no detectable reporter activity after enrichment with FACS. **c,** Flow-cytometry assisted quantification of % GFP positive ABE reporter cells after >10 days of selection for ABE8e-Puro+ cells. Each symbol represents a technical replicate (n=3). Error bars represent mean ± s.d. **d,** Immunoblot analysis for Cas9 expression in B-ALL cells, or 293T cells transfected with ABE8e Puro construct.

**Extended Data Figure 3.**
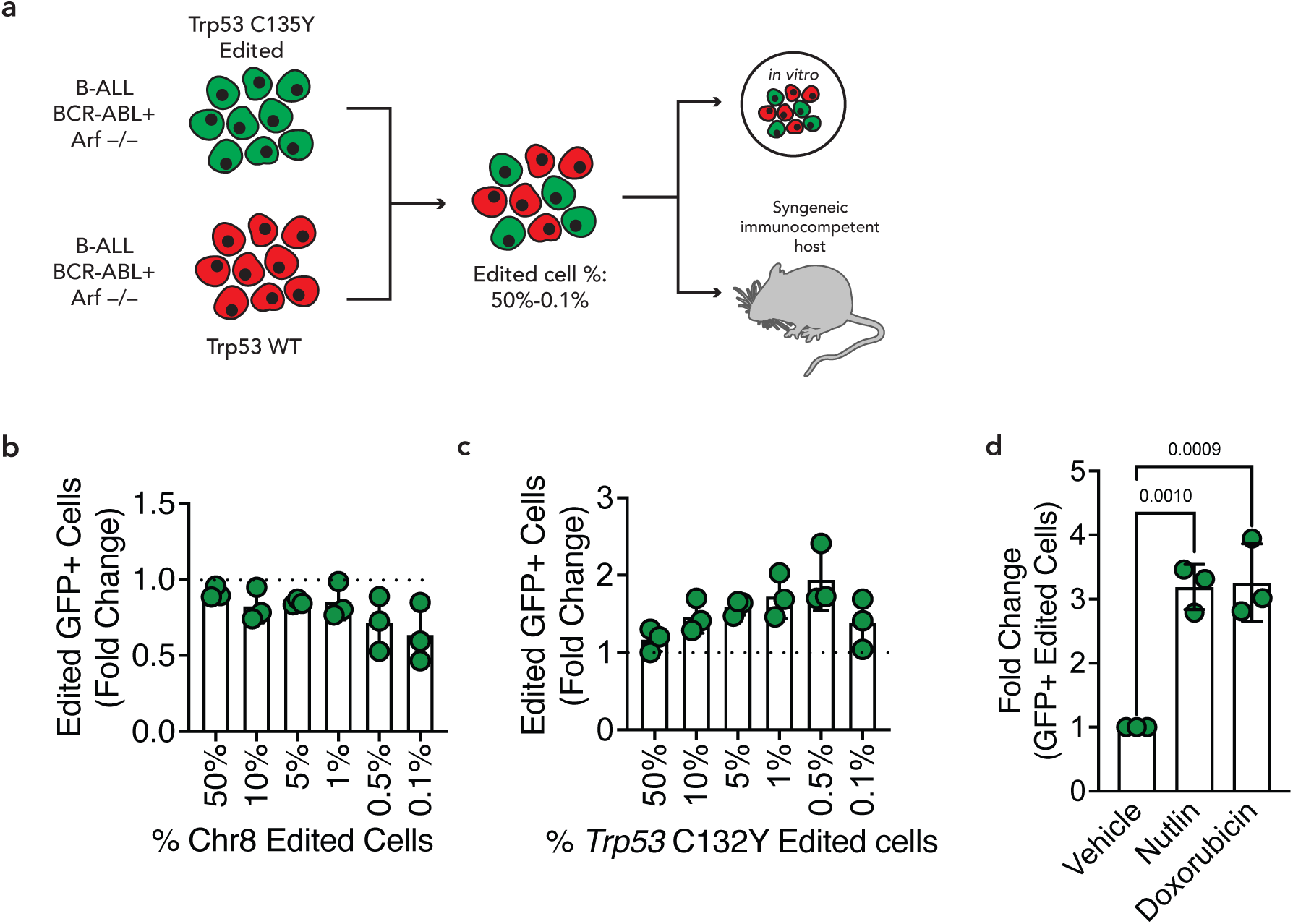
Validation of base editing reporter B-ALL platform. **a,** Schematic of *in vitro* and *in vivo* competition assays between edited and WT B-ALL cells. Distinct percentages of GFP+ *Chr8* (**b**) *or Trp53* (**c**) edited cells were mixed with unlabeled controls at t=0 and differences in growth were measured after 10 days by quantifying the fold-change of GFP+ cells using flow cytometry. Each symbol represents the mean of *n* = 3 technical replicates from n=3 independent experiments. Error bars represent mean ± s.d. Statistical significance determined by two-tailed Student’s *t*-test **d,** Fold change of GFP+ *Trp53* C132Y edited cells after treatment with Nutlin-3a (10uM) or Doxorubicin (80nM) for 48hrs. Each symbol represents the mean of *n* = 3 technical replicates from n=3 independent experiments. Error bars represent mean ± s.d. Statistical significance determined by two-tailed Student’s *t*-test.

**Extended Data Figure 4.**
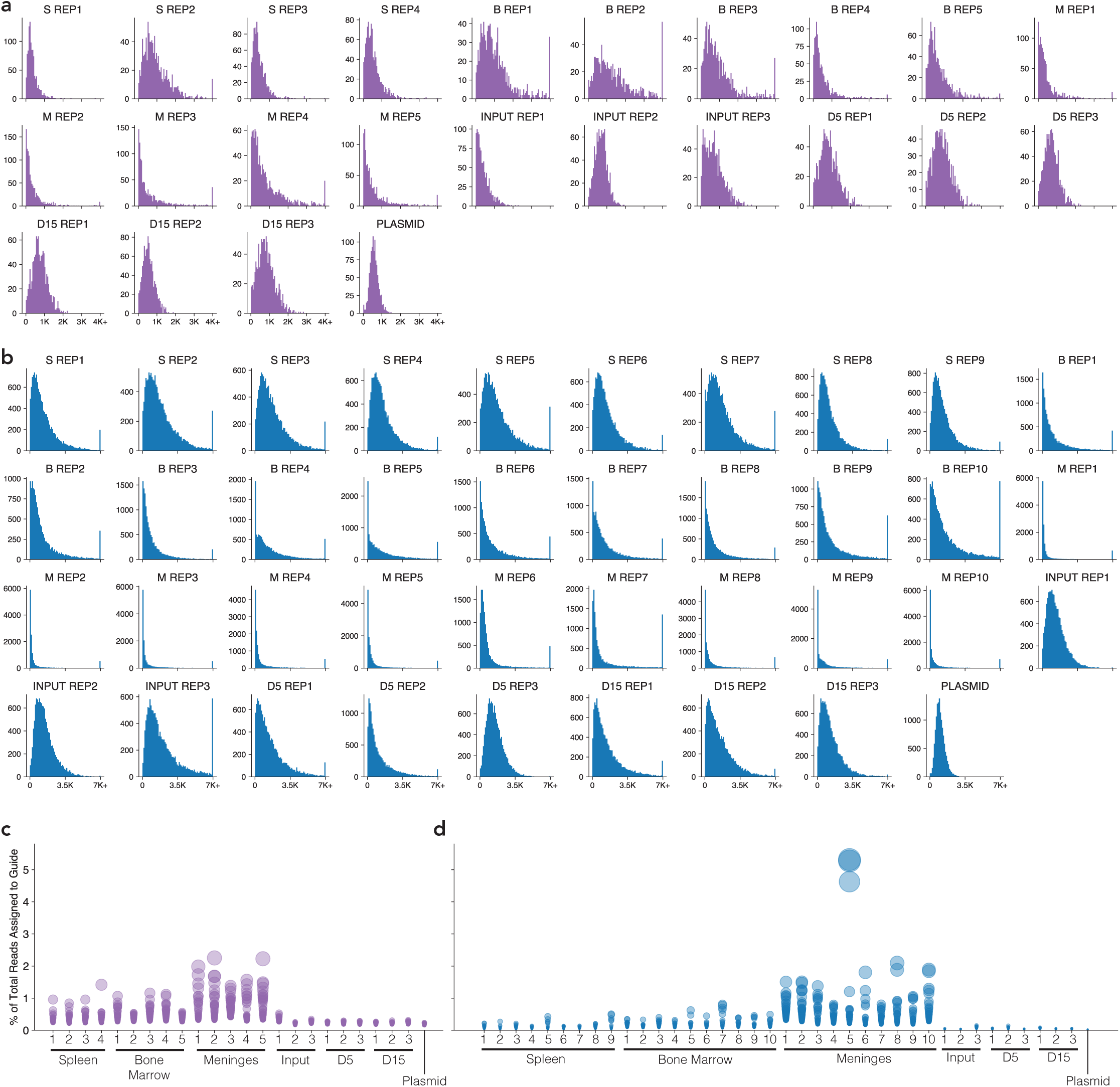
Barcode counts and bottleneck analysis. **a,** Histogram of gRNA counts for individual samples from ABE subpool of MBESv2. S, spleen; B, bone marrow; M, meninges. **b,** Histogram of gRNA counts for individual samples from CBE subpool of MBESv2. S, spleen; B, bone marrow; M, meninges. **c,** Bubble plot of the percentage of total sequencing reads assigned to a given gRNA for the top 50 represented gRNAs in each sample in the ABE subpool. **d,** Same as **(c)**, but for the CBE subpool.

**Extended Data Figure 5.**
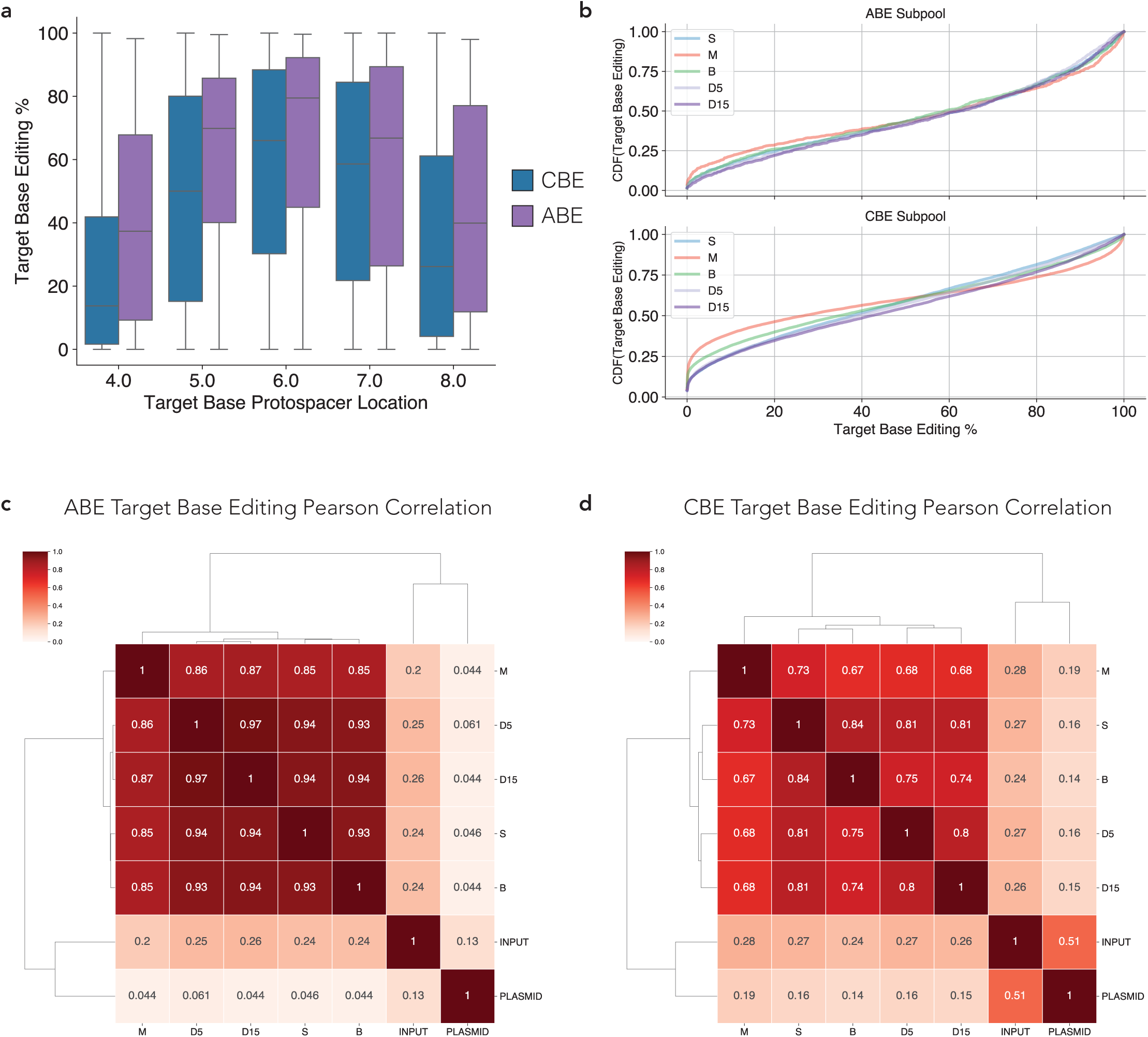
Analysis of sensor editing. **a,** Boxplot of target base editing rate, separated by the protospacer location (+4 to +8) of the target base, for ABE and CBE gRNAs with at least 10 sensor reads. **b,** CDF of target base editing rate for ABE (top) and CBE (bottom) gRNAs, for gRNAs with at least 10 sensor reads. **c,** Correlation matrix of target base editing rate among different ABE samples. **d,** Correlation matrix of target base editing rate among different CBE samples.

**Extended Data Figure 6.**
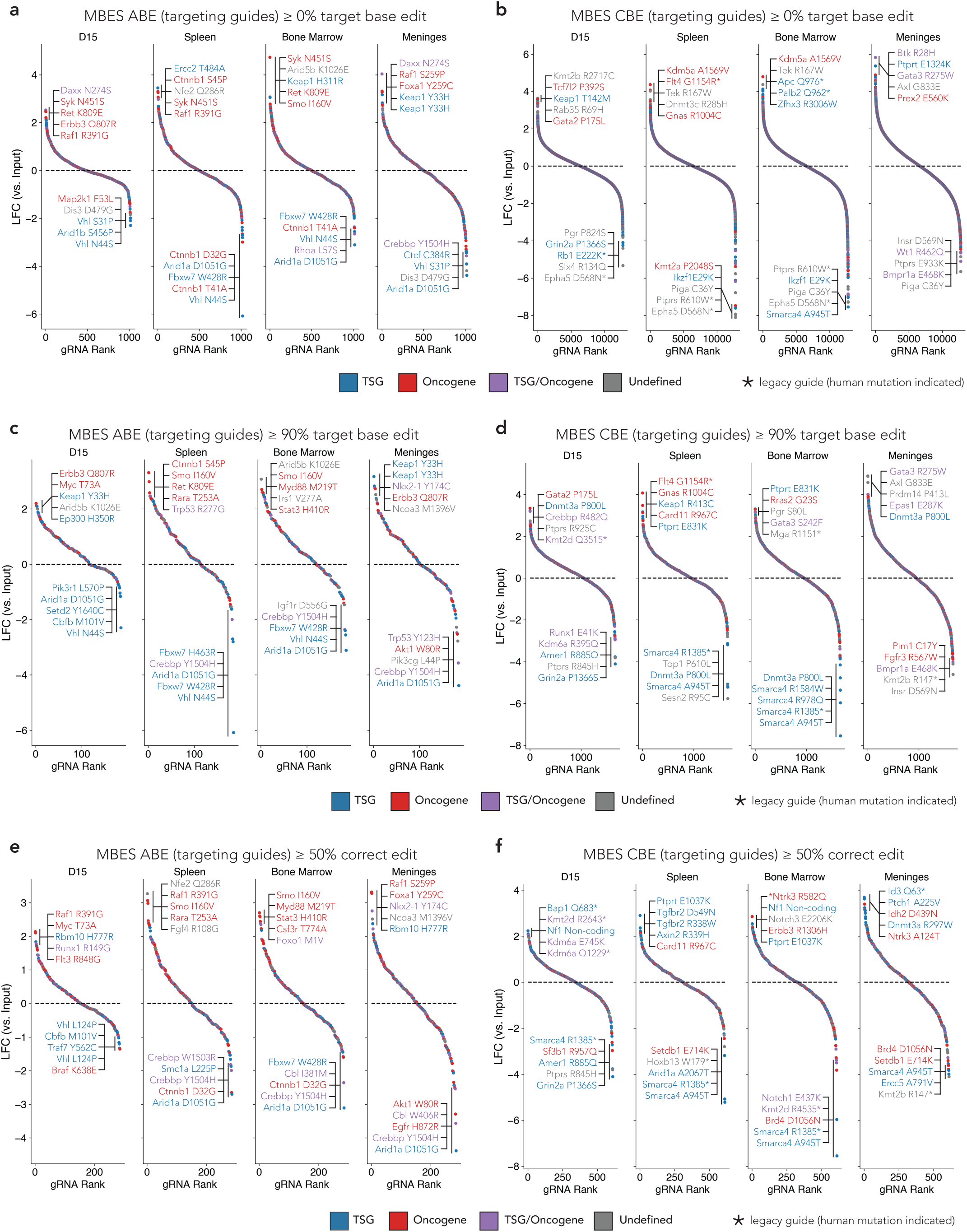
Representative waterfall plots using different sensor editing cutoffs. **a-b,** Waterfall plots of ABE **(a)** or CBE **(b)** gRNA enrichment in each sample type, with top 5 and bottom 5 gRNAs labeled. Guides with fewer than 50 average control counts excluded. Points colored by COSMIC Cancer Gene Census annotation. **c-d,** Same as **(a)** and **(b)** but for gRNAs with target base editing ≥ 90%. **d,** Same as **(b)** but for gRNAs with target base editing ≥ 90%. **e-f,** Same as **(a)** and **(b)** but for gRNAs with correct editing ≥ 50%.

**Extended Data Figure 7.**
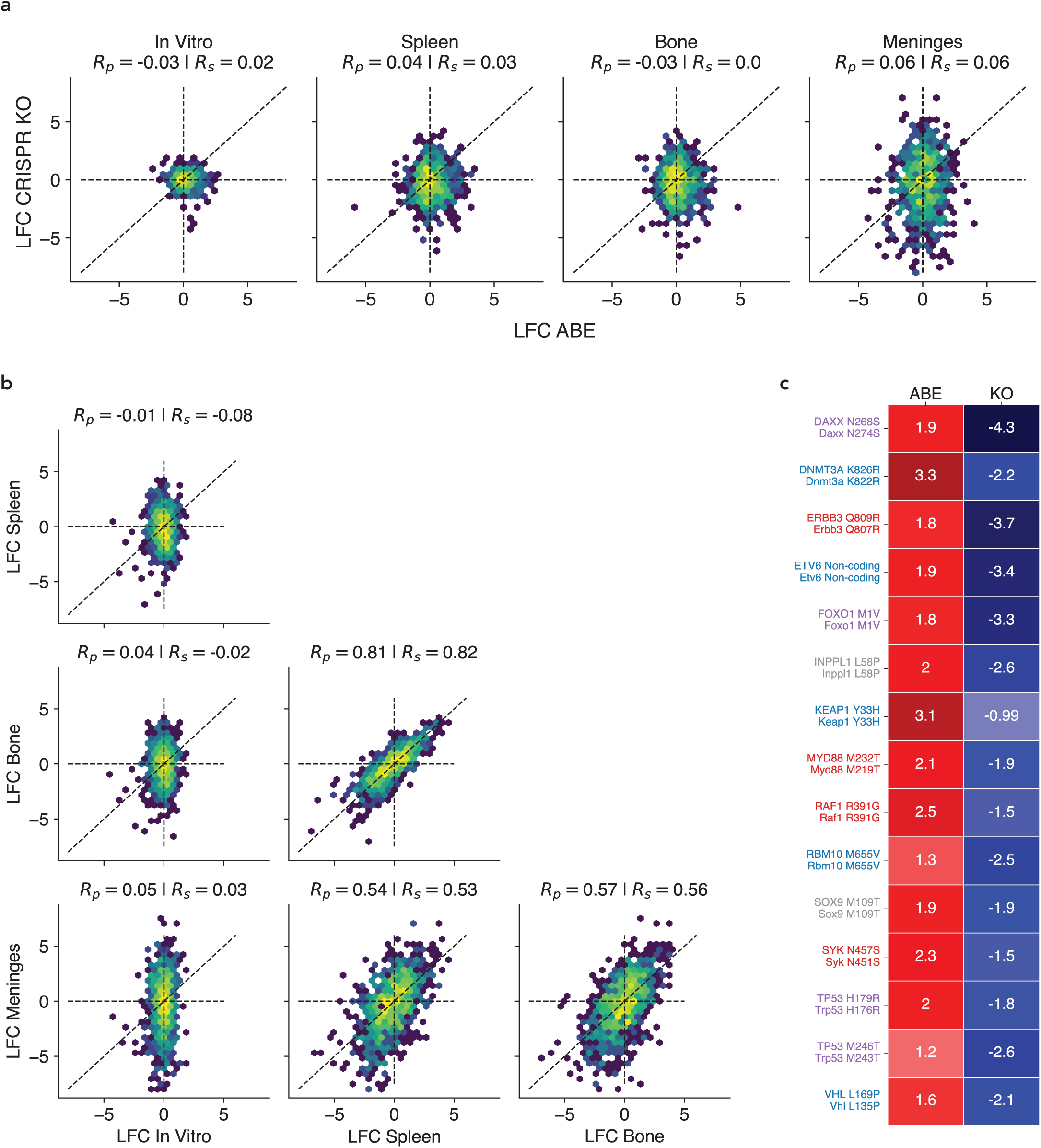
Comparative analysis of CRISPR nuclease and base editing screens. **a,** Scatterplots of gRNA LFC values obtained from CRISPR nuclease (KO) and ABE screens across each context. **b,** Scatterplots of gRNA LFC values obtained from CRISPR nuclease (KO) screens across each context. **c,** Heatmap showing representative gRNAs exhibiting the largest differences in LFC between ABE and CRISPR KO screens. Limited to guides with target base edit ≥ 20%.

**Extended Data Figure 8.**
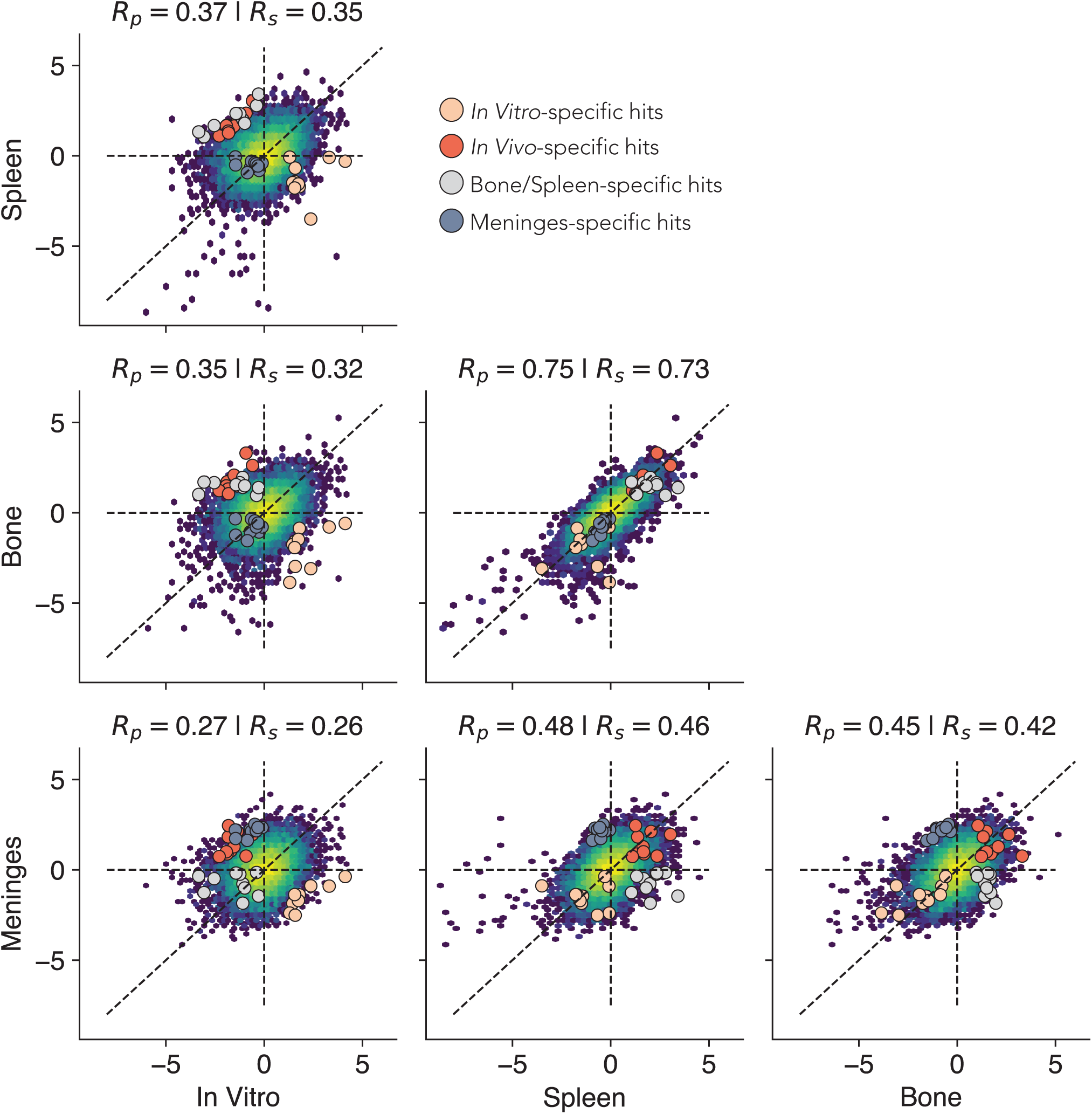
Comparative analysis of base editing screens across experimental contexts. Scatterplots of gRNA LFC values obtained from base editing screens across each context, highlighting putative context-specific mutations following the categories denoted in the sample legend.

## Methods

### Ethics statement

The research conducted in this study was conducted ethically and complies with all relevant guidelines and regulations. Animal studies were performed under strict compliance with the Committee of Animal Care (CAC) at the Massachusetts Institute of Technology (#052102824).

### Animal housing and maintenance

All mouse experiments were conducted under Institutional Animal Care and Use Committee (IACUC)-approved animal protocols at the Massachusetts Institute of Technology (MIT). The mouse strains used in this study included male C57BL/6J mice (The Jackson Laboratories), as indicated in the figure legends. All experimental mice used were 8-12 weeks old. Mice were housed under social conditions (two to five mice per cage) on a 12-hour dark/12-hour light cycle, ambient temperature 21 °C ± 1 °C, and humidity 50% ± 10%. All animals were housed in the pathogen-free animal facility of the MIT Koch Institute, in accordance with the animal care standards of the institutions. Food and water was provided *ad libitum*. All animal research at MIT is conducted under humane conditions with utmost regard for animal welfare. The animal care facility staff is headed by a chief veterinarian and includes a veterinary assistant, animal care technicians, and administrative support. MIT adheres to institutional standards for the humane use and care of animals, which have been established to assure compliance with all applicable federal and state regulations for the purchase, transportation, housing, and research use of animals.

### Animal studies

All mouse experiments were approved by the MIT Committee on Animal Care (CAC) and the Department of Comparative Medicine (DCM). For in vivo screens and survival experiments, B-ALL cells were prepared for transplantation by resuspension in 100-200uL PBS (Corning, 21-031-CV) and loaded in 27.5-gauge syringes (Becton Dickinson, Catalog# BD 305620). All cell solutions were administered by tail vein injection. C57BL/6J mice were injected intravenously with 2.0×10^6^ cells, as indicated in the figure legends. All mice were closely monitored and euthanized upon the appearance of morbidity, in accordance with CAC and DCM policy. The size of each animal cohort was determined by estimating biologically relevant effect sizes between conditions and using the minimum number of animals that could reveal statistical significance using the indicated tests of significance. For in vivo screens, the size of the library utilized also dictated the size of the animal cohort required to achieve at least 1000x representation across replicates.

### Base Editor and Reporter Plasmids

All plasmids were generated using Gibson Assembly strategies using NEBuilder HiFi DNA Assembly Master Mix (NEB cat. no. E2621) following the manufacturer’s protocol. All new plasmids, along with detailed maps and sequences, will be made available through Addgene. The lentiviral cytosine base editor (CBE) reporter (Addgene, cat. no. 136895) was kindly provided by Dr. Lukas Dow. The lentiviral adenine base editor (ABE) reporter was generated through Gibson assembly of a U6-sgRNA-SFFV-GFPAdGO2 gene block (GO2 sequence based on ref. 85) and an IRES-mScarlet fragment into the higher titer pLV backbone^132^. The lentiviral plasmid used to clone and express base editing sensor libraries was assembled by transferring the U6-sgRNA-EFS-Blast-P2A-TurboBFP cassette from pUSEBB^10^ into the higher titer pLV backbone^132^. The CBE6-NG *in vitro* transcription (IVT) template was generated using SpCas9-CBE6b-IVT (Addgene, cat. No. 215834) and replacing the Cas9 sequence with one derived from pLenti-FNLSNG-P2A-Puro (Addgene, cat. No. 136900). The ABE8e-NG IVT template was derived from a pUC19 backbone with a (1) T7 promoter modified to be compatible with the CleanCap co-transcriptional mRNA capping strategy, (2) human alpha-globin derived 5’ UTR, (3) AES-mtRNR1 fusion 3’ UTR, (4) P2A-mScarlet reporter, (5) 120 nucleotide polyA sequence with an intervening ‘G’ nucleotide to minimize recombination, and (6) type II restriction site directly downstream of the polyA to create a clean polyA-terminated linear template. ABE8e (Addgene, cat. no. 138495) was subsequently cloned into this backbone and the Cas9 sequence was replaced with one derived from pLenti-FNLSNG-P2A-Puro (Addgene, cat. No. 136900).

### sgRNA plasmids

We cloned *Esp*3I/*Bsm*BI-compatible annealed and phosphorylated oligos encoding sgRNAs into *Esp*3I/*Bsm*BI-linearized pUSEBB using high-concentration T4 DNA ligase (NEB). A 5′ G (to boost U6 transcriptional initiation) was added to sgRNAs that lacked it either by appending it to the 5′ or by substituting the first nucleotide in the 5′ position for a G. All sgRNA sequences are provided in Supplementary Table 1.

### Virus production

Lentiviruses were produced by co-transfection of HEK293T cells with the relevant lentiviral transfer vector and packaging vectors psPAX2 (Addgene, cat. no. 12260) and pMD2.G (Addgene, cat. no. 12259) using Lipofectamine 2000 (Invitrogen, cat. no. 11668030). Viral supernatants were collected at 48- and 72-h post transfection and stored at −80 °C.

### *In vitro* transcription

The IVT ABE template was linearized with BbsI and purified via phenol chloroform extraction and ethanol precipitation. The CBE IVT template was linearized using a PCR protocol detailed in Doman et al^133^ and purified using the QIAquick PCR purification kit (Qiagen cat. no. 28104) following the manufacturer’s protocol. All IVT reactions were performed on the linearized template for 2 hours @ 37 °C using N1-methylpseudouridine-5’-triphosphate and cleanCap AG co-transcriptional capping (TriLink cat. no. N-1081-10 and N-7113-10). DNA template was then digested away with TURBO DNase (Thermo cat. no. EN0521). mRNA was then purified using lithium chloride precipitation. RNA was quantified and QC was performed by running the RNA on a glyoxal gel to confirm the appropriate size and lack of significant RNA degradation (NorthernMax AM8678). Base editor mRNA was also tested for functionality by nucleofection into reporter cell lines. See Supplementary Protocol 1 for a detailed protocol. Cas9-NG mRNA was generated by TriLink via custom order.

### Immunoblot analysis

Cells were lysed in RIPA buffer, resolved on NuPage 4–12% Bis-Tris protein gels (Thermo Fisher) and transferred to polyvinylidene fluoride (PVDF) membranes. Blocking, primary and secondary antibody incubations were performed in Tris-buffered saline (TBS) with 0.1% Tween-20. Cas9 (1:1000, Cell Signaling, 19526S). Hsp90 was used as a loading control. Protein concentration was determined using a BCA protein assay kit (Pierce).

### Nucleofection

All electroporations were performed using a Neon Transfection System (Invitrogen, discontinued) or a Neon NxT Electroporation System (Invitrogen, cat. no. NEON1SK). For all screens and reporter activity assays, 500,000 cells were electroporated per reaction with the following parameters: 1600V, 20 ms, 1 pulse. Electroporations used either 10 ug (Cas9, ABE) or 20 ug (CBE) of mRNA per reaction.

### Base editor reporter activity assay

To determine optimal mRNA concentrations for screen electroporations, ABE and CBE reporter B-ALL cells were nucleofected with increasing amounts of their respective editor mRNA using the electroporation parameters described above. After 72 hours of recovery post-electroporation, cell fluorescence was measured via flow cytometry, with mScarlet serving as a marker for reporter transduction and GFP indicating successful base editing. Editing efficiency of the bulk population was determined by dividing the number of GFP+, mScarlet+ cells by the number of mScarlet+ cells. Given that they constitutively express an GFP-targeting sgRNA, double positive CBE reporter cells were subsequently used as a reporter for Cas9 activity. Following the same timeline as the base editor activity assay, nuclease efficiency within the bulk population was measured by dividing the number of mScarlet+ cells by the number of mScarlet+, GFP+ cells.

### Cell Lines

All cell lines were mycoplasma negative. Murine *Bcr-Abl*-driven mouse acute B-cell lymphoblastic leukemia cells (Williams et. al. 2006) were cultured in RPMI with L-glutamine (Corning, 10-040-CM), supplemented with 10% fetal bovine serum (FBS), GlutaMAX (Gibco), and 2-mercaptoethanol to a final concentration of 0.05 mM (Gibco, 21985023). B-ALL reporter cell lines were generated by transduction with lentiviruses containing the ABE and CBE reporter constructs described above and grown in the same conditions as *parental* cells. Human HEK293T (ATCC CRL-3216) cells were cultured in DMEM (Gibco) supplemented with 10% fetal bovine serum (FBS) and GlutaMAX (Gibco).

### Flow Cytometry

Single cell suspensions were prepared from cells after collection by passing specimens through a 100-μM cell strainer. Cells were resuspended in FACS buffer (PBS 1X + 2.5% fetal bovine serum). For all experiments, flow cytometric analysis was conducted using either BD FACS Symphony A1 or Celesta analyzers. Fluorescence-assisted cell sorting was performed in either BD FACS Aria II or Sony MA900 cell sorters. Data were analyzed using FlowJo v10.10 as indicated in the figure legends.

### Cloning of MBES v2 libraries

A pooled ABE/CBE sgRNA library was ordered from Twist Biosciences. The lyophilized library was resuspended in 100 µl of TE buffer (pH 8.0) and diluted to create 1 ng µl^−1^ stocks. We performed *n* = 5 (ABE) or *n* = 54 (CBE) PCR reactions with NEBNext High-Fidelity 2× PCR Master Mix (cat. no. M0541S) to amplify the library with the following primers at a low cycle count: ABE forward 5′-AGGCACTTGCTCGTACGACG, ABE reverse 5′-TTAAGGTGCCGGGCCCACAT, CBE forward 5′-GTGTAACCCGTAGGGCACCT, CBE reverse 5’-GTCGAAGGACTGCTCTCGAC. These PCR reactions were pooled and purified using the Qiagen PCR purification kit following the manufacturer’s protocols, with 10 µl of 3 M Na acetate pH 5.2 added for every five volumes of PB used per one volume of PCR reaction. The library was digested with *Esp*3I and *EcoRI* (NEB), pooled, and purified. Subsequently, *n* = 12 (ABE) or *n* = 16 (CBE) ligations were performed using 300 ng of digested and dephosphorylated Trono-BB backbone and 3 ng of digested insert with high concentration T4 DNA Ligase (NEB, cat. no. M0202M). The ligation reactions were precipitated using QuantaBio 5PRIME Phase Lock Gel tubes before being resuspended in 3 µl of EB Buffer per four precipitated reactions. These precipitated ligation reactions were electroporated into Lucigen Endura ElectroCompetent cells (cat. no. 60242-2) before being plated on LB-carbenicillin plates and incubated at 37 °C for 16 h. We scraped the plates and collected the bacteria in 250 ml of LB-ampicillin per four plates, before incubating for 2 h at 37 °C, collecting the bacteria by centrifugation, and proceeding to perform a Qiagen Maxiprep, following the manufacturer’s protocol. Once cloned, the libraries were sent for Amplicon-EZ sequencing (Azenta) to assess representation. Both libraries returned with skew ratios^134^ under 5, demonstrating high fidelity library cloning. Lentivirus was generated via the aforementioned protocol, and viral titer was determined through serial dilutions of virus, transductions in 6-well plates with 5 x 10^5^ B-ALL reporter cells, and measurement of the BFP-positive cell fraction at 72-h post-transduction. All sgRNA sequences in the ABE and CBE library can be found in Supplementary Table 2.

### Screening protocol

Reporter B-ALL cells were transduced with ABE/CBE MBES library virus at a low MOI (between 0.3 and 0.5) to ensure a single sgRNA integration event. Each step of the ABE, CBE, and Cas9 screens—from infection to sequencing—were optimized to achieve a minimum representation of 1,000×. For instance, in the CBE screen, we spinfected a total of 40 million cells across 6-well plates using the volume of viral supernatant that would achieve a 30% infection rate (around 13.3 million transduced cells). BFP, mScarlet double-positive (or BFP, mScarlet, GFP triple positive cells for the Cas9 screen) cells were sorted out 72-h post-transduction and expanded to allow adequate numbers for electroporation. Once expanded, cells were electroporated as described above. The number of cells electroporated per screen accounted for the efficiency of each editor (e.g. CBE6-NG showed an average editing efficiency of 50% with the reporter, so 26 million cells were electroporated to maintain adequate representation). In parallel, three non-electroporated, “input” cell pellets were collected. GFP, BFP, and mScarlet triple-positive cells (or BFP, mScarlet double-positive for the Cas9 screen) were sorted out 72-h post-electroporation. The sorted cells were expanded for 48 hours before preparation for *in vivo* and *in vitro* screening. On screening day, three “day 5” pellets were collected, three *in vitro* replicates were plated, and 2 million cells were injected per mouse. A total of 5 mice were injected for the ABE/Cas9 screen (2000x representation/mouse) and 10 mice for the CBE screen (>1000x representation across all mice). The *in vitro* arms were split as needed, ensuring that a minimum 1000x representation was maintained. The screens were concluded when the mice showed high disease burden ten days later. On that day, *in vitro* “day 15” cell pellets from each replicate were collected. Within the *in vivo* arm, the spleen, bone marrow, and meninges were isolated from each mouse. Prior to sorting, each meninges sample was digested in 1 mL of RPMI with 2.5 mg/mL Collagenase D (Roche, cat. no. 11088858001) and 0.1 mg/mL DNase (Roche, cat. no. 10104159001)^135^ at 37°C for 30 minutes. GFP, BFP, and mScarlet triple-positive cells (BFP, mScarlet double-positive for the Cas9 screen) were sorted out from the single cell suspension of each tissue. A minimum of 1 million cells were collected per tissue in the ABE/Cas9 screen (1000x representation per tissue per mouse) and a minimum of 6.5 x 10^5^ cells were collected per tissue in the CBE screen (>500x representation per tissue across all mice). All sorted cells were pelleted and stored at –20°C.

### Genomic DNA extraction

Genomic DNA (gDNA) was extracted from cells using the DNeasy Blood and Tissue Kit (Qiagen) following the manufacturer’s instructions. Cell pellets were processed in parallel and the resulting gDNA was resuspended in 125 μl of 10 mM Tris-Cl; 0.5 mM EDTA; pH 9.0. Samples from corresponding replicates from MBES and HBES screens were pooled at the gDNA level, measured using a NanoDrop One (ThermoFisher). For low concentration samples, a sodium acetate and ethanol precipitation followed by resuspension in 20 uL of 10 mM Tris-Cl; 0.5 mM EDTA; pH 9.0 was performed prior to screen deconvolution.

### NGS sample preparation

We employed a modified two-step PCR version of the protocol published by Doench et al^136^. adapted to our unique library design. Briefly, we performed an initial PCR, whereby the integrated sensor cassettes were amplified from gDNA, followed by a second PCR to append Illumina sequencing adapters on the 5′ and 3′ ends of the amplicon, as well as a unique demultiplexing barcode on the 5′ end. Each ‘PCR1’ reaction contained 25 μl of Q5 High-Fidelity 2× Master Mix (NEB), 2.5 μl of Sensor_PCR1_F Primer (10 μM), 2.5 μl of Sensor_PCR1_R Primer (10 μM), and up 5 μg of gDNA in 20 μl of water (for a total volume of 50 μl per reaction). PCR1 amplicons were size-selected on a 1% agarose gel, purified using the QIAquick Gel Extraction Kit (Qiagen), and used as template for ‘PCR2’ reactions. Each PCR2 reaction contained 25 μl of NEBNext 2× Master Mix (NEB), 2.5 μl of a common Sensor_PCR2_F (10 μM), 2.5 μl of a uniquely barcoded Sensor_PCR2_R_BARCODE Primer (10 μM), and 10 ng of PCR1 product in 20 μl of water (for a total volume of 50 μl per reaction). We performed one PCR2 reaction per PCR1 product. Library amplicons were size-selected on a 1% agarose gel and purified using the QIAquick Gel Extraction Kit (Qiagen). All primer sequences are available in Supplementary Table 1. PCR program for PCR1 was: (1) 98°C for 30 s; (2) 98°C for 10 s; (3) 64°C for 30 s; (4) 72°C for 30 s; (5) Go to step 2 for 24 cycles; (6) 72°C for 2 min; (7) 4°C forever. PCR program for PCR2 was: (1) 98°C for 30 s; (2) 98°C for 10 s; (3) 70°C for 30 s; (4) 72°C for 30 s; (5) Go to step 2 for 9 cycles; (6) 72°C for 2 min; (7) 4°C forever.

### Next-generation sequencing

For NGS of the CBE screen, we used the NovaSeq SP 300 sequencing system (NovaSeq 6000). NGS for the ABE/Cas9 screen used the MiSeq v3 150 nt sequencing system. Both sequencing runs used a custom sequencing primer set to amplify the protospacer, sensor sequence, and sample barcode in separate reads. All sequencing primers are listed in Supplementary Table 1. The custom sequencing approach for both screens is diagrammed in Supplementary Protocol 2.

### Analytic/computational methods

To analyze NGS data, we used a custom sequencing pipeline to count barcode and protospacer reads, split sensor reads into separate fastq files, and analyze these sensor reads for editing with crispresso2^137^. All data was processed on the Koch Institute’s Luria computing cluster. Code for running this analysis pipeline is available on GitHub: https://github.com/samgould2/base-editing-sensor-analysis.

For the analysis of log2 fold-changes for all samples, we used MAGeCK^138^ on the barcode counts of the gRNAs present in our library. All samples were compared to the “input” barcode counts for the calculation of LFCs. Further, we excluded gRNAs with a control count of <50 normalized barcode counts.

## Data availability

All processed datasets and corresponding source data are available at the following GitHub repository: https://github.com/samgould2/B-ALL-in-vivo-base-editing/.

## Code Availability

Code for running the custom base editing sensor analysis pipeline is available at the following GitHub repository: https://github.com/samgould2/base-editing-sensor-analysis.

All analysis scripts, as well as Jupyter notebooks for generating each figure that appears in the paper, are available at the following GitHub repository: https://github.com/samgould2/B-ALL-in-vivo-base-editing/.

## Data Tables

**Supplementary Table 1 |** Sequences of sgRNAs and primers used in this study.

**Supplementary Table 2 |** MBES v2 library sequences.

**Supplementary Protocol 1 |** *In vitro* mRNA transcription of base editors

**Supplementary Protocol 2 |** Preparing sgRNA-sensor libraries for next generation sequencing Other information and source data are provided in the Github Repository.

## Acknowledgements

We thank Alyna Katti and Lukas Dow for sharing GO base editing reporter plasmids, Emily Zhang and David Liu for sharing SpCas9-CBE6-NGG plasmids, Sujatha Jagannathan for scientific discussions related to her recent discovery of the Gly-PTC mRNA context in triggering NMD (ref. 93), and Yadira Soto-Feliciano for comments and suggestions. We thank Claire Glickman for laboratory management support, Jamie Rothman for administrative support, and members of the Sánchez-Rivera and Hemann laboratories for technical support. We also thank the Koch Institute Swanson Biotechnology Center for technical support, especially the Flow Cytometry Core and the Barbara K. Ostrom (1978) Bioinformatics Facility and the Genomics Facility. Work in the Sánchez-Rivera laboratory is supported by the Howard Hughes Medical Institute (Hanna Gray Fellowship, GT15656), the V Foundation for Cancer Research (V2022-028), NCI Cancer Center Support Grant P30-CA014051, the Virginia and D.K. Ludwig Fund for Cancer Research, Koch Institute Frontier Research Program, the Casey and Family Foundation Cancer Research Fund, the Michael (1957) and Inara Erdei Fund, the MIT Research Support Committee, the Upstage Lung Cancer Foundation, and a Traditional Project Award from the Bridge Project, a partnership between the Koch Institute for Integrative Cancer Research at MIT and the Dana-Farber/Harvard Cancer Center. Work in the Hemann laboratory is supported by the MIT Center for Precision Cancer Medicine, NCI R01-CA233477, R01-CA226898, and NIH/NIAID R21AI151827, as well as NCI Cancer Center Support Grant P30-CA014051 and the Virginia and D.K. Ludwig Fund for Cancer Research. G.A.J., S.I.G., Y.L., D.D., O.A., S.G., and M.E.C. were supported by NIH T32GM136540. G.A.J. was also supported by a Margaret A. Cunningham Immune Mechanisms in Cancer Research Fellowship Award. S.I.G. was also supported by a MIT School of Science Fellowship in Cancer Research. D.D. was also supported by a MIT Dean of Science Fellowship. O.A. was supported by a David H. Koch Graduate Fellowship. V.K.N. was supported by a K12 Paul Calabresi Career Development Award from the NIH/NCI and a Myelodysplastic Syndromes Young Investigator Award from the Edward P. Evans Foundation.

## Author contributions

J.A., G.A.J., S.I.G., M.T.H., and F.J.S.R. conceived the project and wrote the manuscript. J.A. and G.A.J. performed all experiments, with assistance from Y.L., S.G., M.E.C., and A.N.W. S.I.G. performed all computational analyses and assembled all figures. K.D. developed H2M and designed MBESv2 libraries. D.D. and O.A. cloned MBESv2 libraries. J.B. cloned CBE6 and ABE8e NG base editors. V.K.N. generated IVT backbones. M.T.H. and F.J.S.R. supervised the work and secured funding.

## Ethics declarations

F.J.S.R. has consulted for Repare Therapeutics and Ono Pharma. The remaining authors declare no competing interests.

