## Supplementary Protocol 1 for "Multiplexed *in vivo* base editing identifies functional gene-variant-context interactions"

### In vitro mRNA transcription (e.g. for mRNA encoding genome editors)

**Note:** this protocol was developed and optimized by Varun Narendra. Modifications are noted below.

#### Workflow

I - DNA template preparation → II - In vitro transcription reaction → III - DNase treatment → IV - RNA purification → V - Phosphatase treatment → VI - Quality control (QC)

Materials needed:

| | |
|---|---|
| Materials/Equipment | ● |
| 1M Tris pH 8.0 | ● |
| 1M Mg Acetate | ● |
| 7.5M LiCl | ● |
| 1M DTT | ● |
| RNAse free water (Can use MilliQ water) | ● |
| 10% TritonX-100 (DNAse, RNAse free, Fischer, AC327371000) | ● |
| 1M spermidine (Spermidine, free base, 99.5%, MP Biomedicals, ICN15206801, kept @ -20C, soluble in H2O) | ● |
| 100uM N1-methlypseudouridine (N-1081-10) | ● |
| 100uM CleanCap AG (TriLink, N-7113-10) | ● |
| Yeast Inorganic Phosphatase (YIPP, NEB, M2403L) | ● |
| T7 RNA polymerase (NEB, M0251L) | ● |
| NTP Set, 100 mM Solution (Thermo, R0481) | ● |
| Murine RNAse inhibitor (NEB, M0314L) | ● |
| DNase I reaction buffer (B0303S | ● |
| DNase I 1U/uL (Thermo, EN0521) | ● |
| Millennium™ RNA Markers-Formamide (Thermo, AM7151) | ● |
| NorthernMax-Glyoxal Loading Dye (AM8551) | ● |
| NorthernMax Running Buffer (AM8678) | ● |

| | |
|---|---|
| Ambion PureLink RNA mini kit (Thermo, 12183018A) | • |

#### Considerations

- Note: RNA is very sensitive to degradation by RNAses.
- To minimize RNA degradation:
    - Clean the bench and all tube racks w/ 10% bleach → RNAse-ZAP → 70% Ethanol.
    - Clean all pipettes and pipet aid w/ RNAse-ZAP → 70% Ethanol
    - Spray everything else thoroughly with RNAse-ZAP → 70% Ethanol
    - Ensure all buffers and water used are clean and RNAse free.
    - Ensure all tips used are filter tips and are RNAse free.
    - Ensure that the purified plasmid DNA is RNAse free. These preps are ideally stored in DNA LoBind tubes that are exclusively used for RNA work. If you are not sure that the plasmid prep is RNAse free, re-prep or extract w/ equilibrated phenol.
    - Use DNA LoBind tubes for all steps.
- Fast-temp a centrifuge prior to starting precipitation steps.
- Prepare 100% and 70% RNAse free ethanol in 50mL conical tubes. Leave at -20C.
- Carry out all incubations in a thermal cycler.
- Pellets from the digested IVT template plasmid may be really difficult to see since the amount of starting DNA material is low. Therefore, centrifuge all tubes in the same orientation such that you know exactly where the pellet will be. You can then aspirate the liquid from the opposite side of the tube.
- When using Ambion PureLink columns, do a total of three separate elutions per column and store them separately. Measure all elutions and pool the ones that are well concentrated (>1-1.5ug/uL). The reason for doing this is that these columns still have a good amount of RNA bound to them after the first and second elutions.
- Always QC the RNA using glyoxal gels to confirm the mRNA is of the correct size. Make sure you have the appropriate running buffers, loading dye, and millennium marker (can get all of these from Thermo). These gels can be run at 150-200V.

### Protocol

##### I - DNA template preparation

For ABE templates (cloned into pUC19 backbone):

1. Digest backbone w/ BbsI-HF (set these up in RNase-free PCR strips):
    a. 50uL reaction:
        i. 50ug Template plasmid DNA (X uL)
        ii. H2O (up to 40uL)
        iii. 10X cutsmart buffer (5uL)
        iv. BbsI-HF (20U/ul) (5uL)
        v. Incubate @37C for 5hrs in thermal cycler
2. Phenol-chloroform precipitation to purify:

a. Pre-spin PhaseLock tubes at 13K RPM for 5 minutes.
b. Transfer digestion to DNA LoBind 1.5ml Eppendorf tubes.
c. Raise reaction to 500uL with H2O.
d. Add 500uL Phenol:Chloroform:Isoamyl alcohol. Vortex for 10-15''.
e. Transfer solution to PhaseLock tube.
f. Spin at 13K RPM x 5'.
g. Transfer 400uL of aqueous phase to a new LoBind tube. Avoid any phenol contamination as this will inhibit downstream IVT reaction.
h. Add 44uL 3M Na Acetate pH 5.2. Mix well by flicking tube.
i. Add 1mL ice-cold 100% ethanol
j. Incubate @ -80C for minimum of 30'. Yield will increase with longer incubations (i.e. O/N @ -20C)
k. Spin at 20k x g for 30' @ 4C (fast-temp centrifuge prior).
l. NOTE: Pellet may not be clearly visible w/o addition of glycogen. If not visible, avoid aspirating to bottom of the Eppendorf tube to prevent aspiration of the pellet.
m. Wash x2 with ice-cold 70% ethanol with 5' max-speed spins after each wash.
n. After last wash, spin down 20k x g for 1' to recover all residual ethanol at bottom of tube. Remove remaining ethanol with 10uL pipette tip w/o disturbing pellet.
o. Resuspend in 20-30uL H2O, depending on the size of the pellet. Quantify on nanodrop.

For CBE6 IVT templates (CBE6 originally described in PMID: 38402281; additional details discussed in PMID: 35941224)

1. DNA templates for in vitro transcription must be linear not circular. To generate a linear in vitro transcription template, PCR amplify from mRNA transcription template plasmids using the primers listed in the table below. Set up the following reaction:

| Component | Amount |
|---|---|
| In vitro transcription forward primer, 100 µM: TCGAGCTCGGTACCTAATACGACTCACTATAAGG | 0.75 uL |
| In vitro transcription reverse primer, 100 µM: TTTTTTTTTTTTTTTTTTTTTTTTTTTTTTTTTTTTTTTTTTTTTTTTTTTTTTTTTTTTTTTTTTTTTTTTTTTTTTTTTTTTTTTTTTTTTTTTTTTTTTTTTTTTTTTTTTTTTTTCTTCCTACTCAGGCTTTATTCAAAGACCA | 0.75 uL |
| mRNA transcription template plasmid, 40 ng µL−1 | 6 uL |
| Phusion U green multiplex master mix, 2× | 150 uL |

| | |
|---|---|
| Nuclease-free H2O | 142.5 uL |
| Total reaction volume | 300 uL |

This reaction is a scaled-up version of a standard 50 μL PCR. We find that total DNA yields from this PCR can be relatively low and that pooling multiple 50 μL PCRs into a single PCR purification column (Step 6) provides enough template for subsequent in vitro transcription (Step 8). Using typical equipment, this 300 μL mastermix will need to be divided into six individual 50 μL reactions on a thermocycler.

2. Perform the PCR under the following conditions: (1) 98°C for 2 mins; (2) 98°C for 15 s; (3) 71.4°C for 30 s; (4) 72°C for 30s/kb (depends on template, use 3 min for CBE6); (5) Go to step 2 for 34 cycles; (6) 72°C for 5 min; (7) 4°C forever.

3. Purify the PCR products from the 300 μL master mix using a single silica column from the QIAquick PCR purification kit (Qiagen) according to the manufacturer's protocols. Elute in EB (provided with the kit) and quantify purified product concentration by UV-visible spectrophotometry (NanoDrop) or equivalent method. Verify product is expected size via 1% TAE gel.

##### II - In vitro transcription reaction

1. Prepare fresh 10X reaction buffer as follows:

| 10X Buffer | [Final] | [Stock] | DF | 5mL 10X Buffer (uL) |
|---|---|---|---|---|
| Tris-HCl pH 8.0 | 400 | 1000 | 2.5 | 2000 |
| Mg Acetate | 165 | 1000 | 6.1 | 825 |
| DTT | 100 | 1000 | 10 | 500 |
| Spermidine | 20 | 1000 | 50 | 100 |
| Triton X-100 (RNAse free) | 0.2% | 10 | 50 | 100 |
| H2O | NA | NA | NA | 1475 |

   a. Store 10X buffer as single-use aliquot at -20C
   b. Note: ensure DTT is 0.2um filtered prior to use and stored at -20C
2. Perform IVT reaction as follows.
   a. Typically, 100uL reaction will generate ~100-200ug of mRNA.

   b. Note: DNA template should be at a final concentration of 25-50ng/uL, so a 50uL reaction will require roughly 1.25 ug digested template.

| IVT reaction | [Final] | 1000uL | 100uL | 50uL |
|---|---|---|---|---|
| Water | - | / | / | / |
| ATP | 5mM | 50 | 5 | 2.5 |
| CTP | 5mM | 50 | 5 | 2.5 |
| GTP | 5mM | 50 | 5 | 2.5 |
| UTP (modified N1-me-PseudoU or methoxy-U) | 5mM | 50 | 5 | 2.5 |
| CleanCapAG | 4mM | 40 | 4 | 2 |
| 10X Buffer | 1X | 100 | 10 | 5 |
| DNA template 25-50 ng/uL | 25-50 ng/uL | X | X | X |
| RNase inhibitor | *1 U/uL* | 25 | 2.5 | 1.25 |
| Yeast inorganic pyro | 0.002 U/uL | 25 | 2.5 | 1.25 |
| T7 RNA polymerase | 8 U/uL | 160 | 16 | 8 |
| Total | | 1000 | 100 | 50 |

3. Incubate for 2 hours at 37C. Hold at 4C.

**III - DNase treatment**

Any residual DNA needs to be removed, as this will be toxic to cells, and affect RNA quantification.

1. Add appropriate volume of 10X DNase I reaction buffer (with MgCl2), and then DNaseI to a final concentration of 0.25 U/ul.
   a. For example, add 15uL of DNaseI buffer and 37.5uL of DNaseI to a 100uL reaction
   b. Do not vortex DNase I (very sensitive to physical denaturation)
   c. Note: 1U degrades 1 ug of DNA in 10’ at 37C.
   d. Reaction requires Mg or Mn. Mg is in the 10X NEB B0303S Rxn buffer. Do NOT use the “w/o Mg” buffer.
2. Incubate @ 37C for 30’.
3. Transfer reaction to a new LoBind tube and proceed directly to the purification step.

**IV - RNA purification**

There are 2 ways of doing this: LiCl precipitation or column purification.

1. **Option 1: LiCl purification**
   a. Raise with H2O to 500uL.
   b. Add 250uL 7.5M LiCl → -20C for 30’.
   c. Spin @ 4C x 30’.
      - Note: Longer -20C incubation (i.e. O/N) will increase yield.
   d. Wash 2X with 70% ethanol.
   e. Spin a final time after 2nd wash to bring residual ethanol to bottom of tube.
   f. Remove any residual Etoh with 10uL pipette, w/o aspirating pellet.
   g. Air dry pellet x 2’.
   h. Resuspend in desired amount of RNase free H2O.
   i. Let it sit at RT for 5-10’ before quantifying on nanodrop. Expected 260:280 ~1.8.
2. **Option 2: Column purification (Ambion)**
   a. Add 10uL of BME (14.3M in hood) to 1mL Lysis buffer.
   b. Raise reaction to 200uL with RNAse free water.
   c. Add 1 volume (200uL) lysis buffer with BME. Pipette u/d 5x.
   d. Add 1 volume (200uL) 100% ethanol. Pipette u/d 5x.
   e. Transfer to Ambion PureLink spin column
   f. Spin 12k x g for 15’’
   g. Add 500uL Wash buffer II (should have EtOh added previously) -> Spin 12k x g for 15’’
   h. Repeat Wash II
   i. Spin 12k x g for 2’
   j. Air dry column for 1’
   k. Add 35uL RNAse free H2O (provided in kit) -> incubate at RT for 2’. Move spin column to a new LoBind tube.
   l. Centrifuge 15k x g for 2’.
   m. Repeat elution (i.e. add 35uL RNAse free H2O) -> elute in same or a different tube, depending on desired RNA concentration.
   n. Quantify on nanodrop. Ensure 260:280 in 1.8-2.0 range. Ideally, it is ~1.8. Ensure 260:230 > 2.

   Note: For elution prior to phosphatase step, can elute in larger volume (i.e. 100uL x 2 elutions)

**V - Phosphatase treatment**

This step will minimize immune activation via 5’ Tri-phosphate recognition on mRNA. 1U dephosphorylates 1ug pUC19 at 37C in 30’. Note, pUC19 is 3kb, and our templates are much larger (~10kb), so we are in effect digesting less molecules for same amount of DNA.

*Note: While we recommend performing this step, we have sometimes skipped it without any major detrimental effects to the functionality of the BE mRNA. Skipping this step would save you from performing a second purification.*

1. Add 10X phosphatase buffer (5.8uL to 50uL reaction)
2. Add 3uL phosphatase (5U/ul) to 50uL reaction.
3. Incubate 37C for 30'.
4. Proceed directly to mRNA purification (as above). Elute/resuspend mRNA in small volume to achieve high concentration mRNA (2-5ug/ul).

**VI - Quality control (QC)**

Before starting, RNAse-zap a gel electrophoresis chamber ahead of running gels, as well as gel comb, gel tray, flask, and thermometer. There are 2 ways of QC'ing using electrophoresis: running a formaldehyde-Agarose gel or running a glyoxal gel.

1. **Option 1: Formaldehyde-Agarose gel:**
    a. To prepare 1% gel, melt 1g of agarose in 88mL H2O. Cool to 60C.
    b. Add 10mL of MOPS buffer.
    c. Add 2.7mL 37% formaldehyde. Poor onto gel apparatus.
    d. Fill gel-tank with 1X MOPS buffer
    e. Load 5uL of ssRNA ladder
    f. Load samples using 2X RNA dye
    g. Heat samples to 70C for 10' and then place directly on ice for 2-5 min prior to loading gel.
    h. Run @ 5-6 V/cm (~75V) until bromophenol blue reaches ~2/3 the length of the gel. This takes ~2-3 hrs.
    i. After the run is complete, stain with SYBR-GOLD (1:10K) in 100mL of TBE, rotating gently. Cover gel with aluminum foil. Ensure the gel is fully submerged.
2. **Option 2: Glyoxal gel:**
    a. Add 10mL of 10X NorthernMax Running buffer to 90mL RNAse free H2O. Transfer to flask.
    b. Add 1g agarose (1% gel)
    c. Heat 1' in microwave.
    d. Cool to 60C and then pour gel.
    e. Dilute 1ug mRNA in total of 5ul H2O. Add 5uL 2X Glyoxal loading dye. In another well, add 5uL millennium RNA marker to 5uL loading dye. Heat at 50C x 30' (can reduce to 15' if in rush). Then transfer to ice directly for 2-5', and load gel. Add 2 drops of EtBr to bottom of gel tank.
    f. Run @5-6V/cm until dye is ½ way through gel.
    g. Image EtBr

Lastly, dilute mRNA to desired concentration using H20, aliquot in RNAse-free PCR strips in single-use aliquots, and store at -80C.

**<u>References</u>**

Ali N, Rampazzo RCP, Costa ADT, Krieger MA. Current Nucleic Acid Extraction Methods and Their Implications to Point-of-Care Diagnostics. Biomed Res Int. 2017;2017:9306564. doi: 10.1155/2017/9306564. Epub 2017 Jul 12. PMID: 28785592; PMCID: PMC5529626.

Doman JL, Sousa AA, Randolph PB, Chen PJ, Liu DR. Designing and executing prime editing experiments in mammalian cells. Nat Protoc. 2022 Nov;17(11):2431-2468. doi: 10.1038/s41596-022-00724-4. Epub 2022 Aug 8. PMID: 35941224; PMCID: PMC9799714.

NEB Labs. (n.d.). DNA Template Preparation (E2040). *neb.com/.* Retrieved from: https://www.neb.com/protocols/0001/01/01/dna-template-preparation-e2040

Zhang E, Neugebauer ME, Krasnow NA, Liu DR. Phage-assisted evolution of highly active cytosine base editors with enhanced selectivity and minimal sequence context preference. Nat Commun. 2024 Feb 24;15(1):1697. doi: 10.1038/s41467-024-45969-7. PMID: 38402281; PMCID: PMC10894238.
