## Supplementary Protocol 1 for "Multiplexed *in vivo* base editing identifies functional gene-variant-context interactions"

### Preparing sgRNA-sensor construct for Next Generation Sequencing (NGS)

- Extract gDNA from 5M cells using “Genomic DNA extraction from cells using the QIAGEN DNeasy Blood & Tissue Kit”.
- For each sample use 10ug of DNA (or divide your total volume of DNA into 4-6 PCR reactions) to perform PCR reactions using Q5 High Fidelity 2X Master Mix (NEB #M0429S).
  - To maintain ≥1000X representation

| **Component** | **50uL Reaction** | **Master Mix (X36) X40** |
|---|---|---|
| Q5 High-Fidelity 2X Master Mix | 25 uL | 1000 uL |
| 10 uM Forward Primer | 2.5 uL | 100 uL |
| 10 uM Reverse Primer | 2.5 uL | 100 uL |
| Template DNA | 20 uL | 20 uL each |
| $H_2O$ | - | |
| **Total** | **50 uL** | **1200 uL** |

For master mix: 30uL to each PCR tube

Trono.BR applicable PCR1 primers (adds the partial illumina adapter)

| **Primer** | **Sequence** |
|---|---|
| Sensor_PCR1_F | CGCTCTTCCGATCTCTAGCGTTCGAGTTAGGAATT |
| Sensor_PCR1_R | CTGAACCGCTCTTCCGATCTTTGTGGAAAGGACGAAACACC |

- Cycle conditions (**x 25 cycles total**):

| Temperature (°C) | cycle no. |
|---|---|
| 98 | 30 s |
| 98 | 10 s |
| 64 | 30 s |

| | |
|---|---|
| 72 | 30 s |
| 72 | 2 min |
| 4 | keep |

- Pool up to 4 PCR reactions and purify using PCR purification kit (QIAquick PCR Purification Kit) according to manufacturer's protocol and elute in 50uL of EB buffer.
- Add 10uL of 6X loading dye to each sample eluted.
- Run samples on a 1% agarose gel alongside a 100bp ladder.
- Gel purify the correctly sized band using (Qiagen Gel Extraction Kit) and elute in 30uL of EB buffer.
- Measure DNA concentration using the NanoDrop 2000 (ThermoFisher).

Sensor-PCR1 product

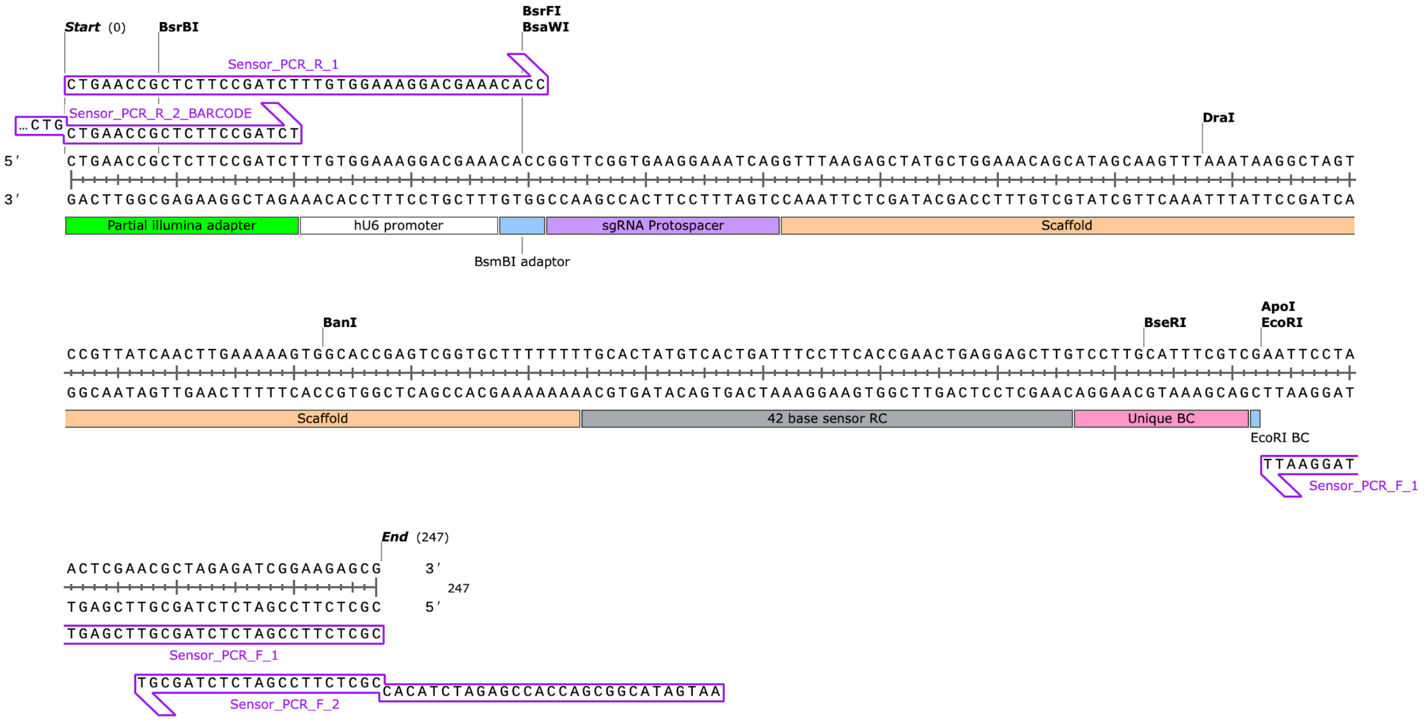

- Use 10ng of PCR1 product to perform 4 PCR reactions using Q5 2X

| Component | 50uL Reaction | Master Mix (X4) X5 |
|---|---|---|
| **Q5 High-Fidelity 2X Master Mix** | 25 uL | 125 uL |
| **10 uM Forward Primer** | 2.5 uL | 12.5 uL |
| **10 uM Reverse BARCODED Primer** | 2.5 uL | 12.5 uL |
| **Template DNA (10ng/uL)** | 1 uL | 1 uL each |
| **H2O** | 19 uL | 95 uL |
| **Total** | **50 uL** | **250 uL** |

Note: make sure you use a different barcoded primer for each sample

PCR2 primers bind the illumina partial adaptors in add unique sample barcodes as well as i7 and i5 anchor sequences required for NGS

| Primer | Sequence |
|---|---|
| Sensor_PCR2_F | AATGATACGGCGACCACCGAGATCTACACCGCTCTTCCGATCTCTAGCGT |
| Sensor_PCR2_R_BARCODE | CAAGCAGAAGACGGCATACGAGATNNNNNNNNCCTGCTGAACCGCTCTTCCGATCT |

For master mix: 49uL to each PCR tube

- Cycle conditions (**x 10 cycles total**):

| Temperature (°C) | cycle no. |
|---|---|
| 98 | 2 min |
| 98 | 10 s |
| 70 | 30 s |
| 72 | 30 s |

| | |
|---|---|
| 72 | 2 min |
| 4 | keep |

- Pool up to 4 PCR reactions and purify using PCR purification kit according to manufacturer's protocol and elute in 50uL of EB buffer.
- Add 10uL of 6X loading dye to each sample eluted.
- Run samples on a 1% agarose gel alongside a 100bp ladder.
- Gel purify the correctly sized band and elute in 30uL of EB buffer.
- Measure DNA concentration using the NanoDrop.

Sensor-PCR2 product

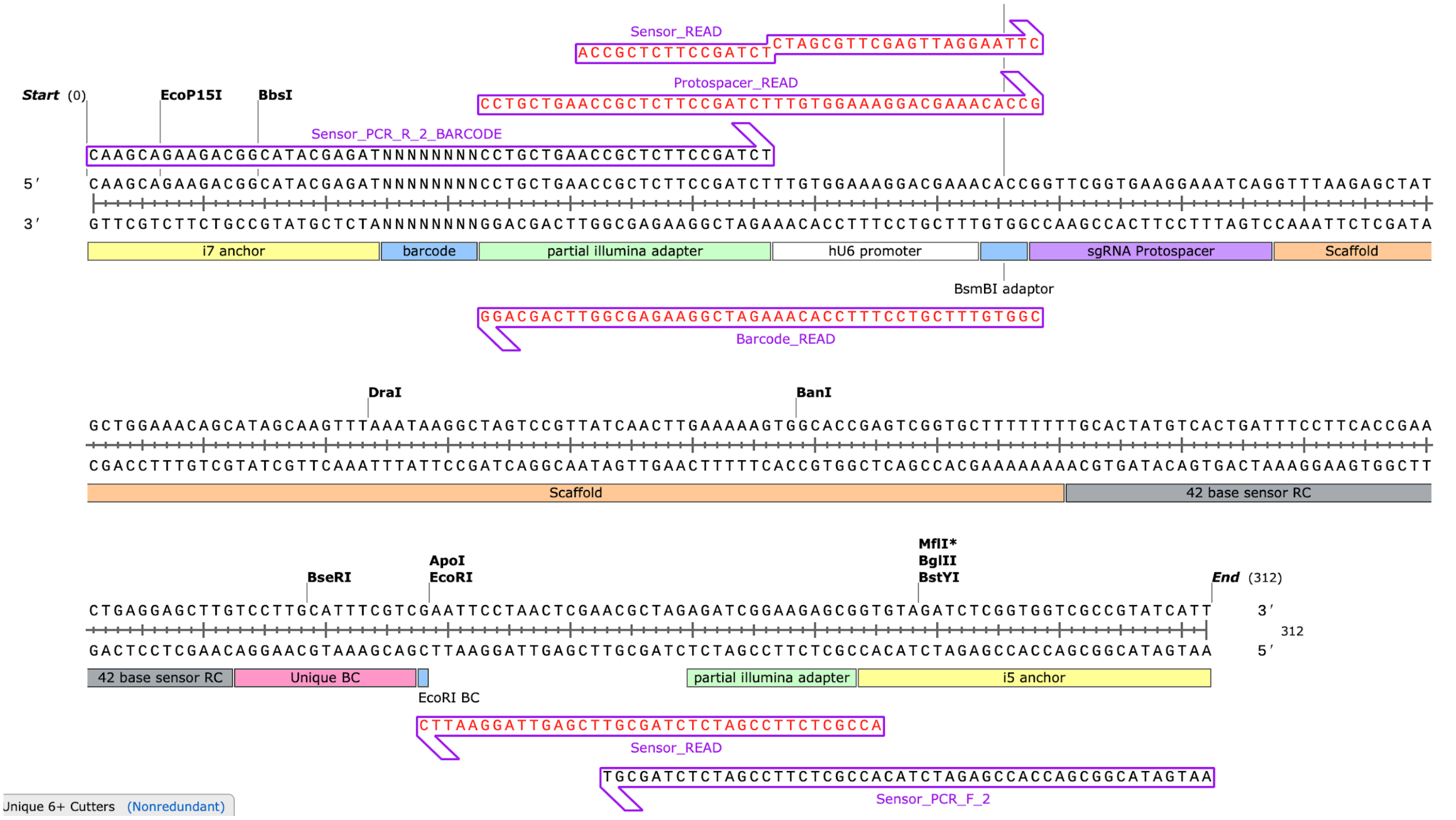

- Submit with the appropriate primers.

Examples:

| Primer | Sequence |
|---|---|
| Protospacer_Read | CCTGCTGAACCGCTCTTCCGATCTTTGTGGAAAGGACGAAACACCG |
| Sensor_Read | ACCGCTCTTCCGATCTCTAGCGTTCGAGTTAGGAATTC |
| Barcode_Read | CGGTGTTTCGTCCTTTCCACAAAGATCGGAAGAGCGGTTCAGCAGG |

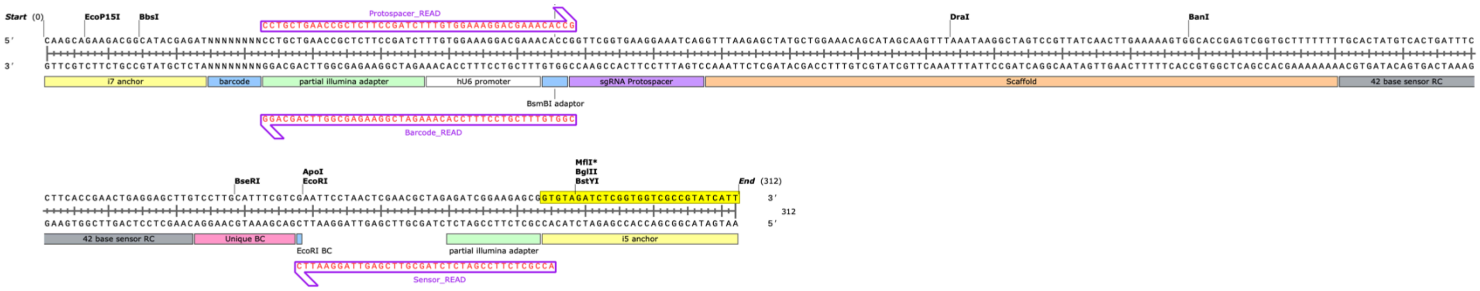
